# Tumor-derived Extracellular Vesicles Induce ER Stress to Drive Tolerogenic Dendritic Cell Development in the Tumor Microenvironment

**DOI:** 10.64898/2026.02.10.705213

**Authors:** Xueying Wang, Michael P. Plebanek, Y-Van Nguyen, Mahere Razazade Bazaz, Michael Sturdivant, Balamayooran Theivanthiran, Brent A. Hanks

## Abstract

**Background:** The efficacy of immune checkpoint blockade relies on the robust priming of T cells by immunostimulatory dendritic cells (DCs). However, the tumor microenvironment (TME) frequently drives DCs into a dysfunctional, pro-tolerogenic state governed by aberrant metabolic rewiring, creating a barrier to durable antitumor immunity. While tumor-derived extracellular vesicles (EVs) are abundant in the TME, their specific role in orchestrating this immunosuppressive metabolic reprogramming remains poorly understood. This study provides insight into the signaling axes through which tumor-derived EVs alter DC function and evaluates the therapeutic potential of targeting these pathways to overcome immunotherapy resistance.

**Methods:** Tumor models were engineered to express EV fluorescent markers to track tumor EV uptake *in vivo*. Bulk and single-cell RNA sequencing was integrated with multi-parameter flow cytometry to characterize the reprogramming of tumor EV-educated DCs both *in vitro* and *in vivo*. Western blotting, quantitative real-time polymerase chain reaction assays, various cellular metabolic assays, as well as T cell-based immunologic studies were utilized to characterize the underlying mechanisms of tumor EV-mediated DC reprogramming. DC-specific *Ppara*-deficient mice were developed to verify these mechanisms *in vivo*. PPAR-α targeted inhibitors were evaluated based on their ability to overcome checkpoint inhibitor resistance in an autochthonous model of melanoma.

**Results:** Tumor-derived EVs were found to promote tumor progression by suppressing host immunity. Further studies reveal that tumor-derived EVs induce a tolerogenic ’mregDC’ transcriptional signature characterized by the upregulation of immunoregulatory molecules in DCs both *in vitro* and *in vivo*. These tumor EV-educated DCs exhibit an impaired capacity for CD8^+^ T cell priming, while demonstrating a proficiency for promoting CD4^+^FoxP3^+^ regulatory T cell differentiation. Mechanistically, tumor EVs concurrently trigger the unfolded protein response (UPR) via the PERK-ATF4 and IRE1α-XBP1s signaling axes, subsequently activating the SREBP2 and PPAR-α transcription factors, respectively. This process drives both aberrant lipid accumulation and fatty acid oxidation (FAO) in DCs residing within the TME. DC-restricted ablation of PPAR-α significantly reversed the pro-tolerogenic effect of tumor EVs *in vivo* while pharmacologic targeting of PPAR-α overcomes anti-PD-1 resistance and augments CD8^+^ T cell infiltration in an autochthonous model of melanoma.

**Conclusions:** Tumor EVs contribute to the development and pro-tolerogenic function of mregDCs in the TME by triggering the UPR pathway. Aberrant lipid metabolism involving enhanced FAO are common characteristics associated with DC dysfunction in the TME. Strategies to interrupt these pathways represent promising approaches for reversing immune tolerance and enhancing tumor-targeted CD8^+^ T cell responses.

## INTRODUCTION

Tumor immunotherapy has revolutionized traditional cancer treatment^1^. However, while immune checkpoint blockade can mediate a long-lasting clinical benefit, only a fraction of patients respond to this therapeutic strategy^2^ ^3^. This is due to the emergence of numerous primary and secondary resistance mechanisms that prevent the efficient activation of immunity^4^ ^5^. Dendritic cells (DCs) have been implicated as a critical mediator of responses to anti-PD-1 immunotherapy and are the most potent antigen-presenting cells due to their ability to efficiently cross-present internalized exogenous antigens on major histocompatibility complex (MHC) class I molecules to stimulate effector CD8^+^ T cells^6–8^. At the early stages of tumor progression, DCs engulf tumor debris and migrate from the tumor bed to tumor-draining lymph nodes (TDLNs), where they present antigens to induce T cell activation and differentiation^9^. Antigen cross-presentation is especially crucial for tumor antigen recognition by T cells as it is indispensable for epitope spreading, a process that drives the stimulation of T cells capable of recognizing a diverse array of tumor antigens from initial dominant epitopes and is highly correlated with the efficacy of immunotherapy^10^ ^11^. To escape immune surveillance, tumors have developed mechanisms that drive DCs to a dysfunctional state in a process denoted as DC tolerization^12–15^.

It has become clear that DC function is highly dependent upon metabolic state^16–19^. We, as well others, have previously demonstrated that tumor cells drive DC tolerization through the induction of fatty acid oxidation (FAO)^20–22^. Indeed, our studies indicate that the recently described ’mature DCs enriched in immunoregulatory molecules’ (mregDC) population represents a conventional DC population that is functionally pro-tolerogenic while also exhibiting both increased lipid stores and elevated levels of FAO^23^ ^24^. The tumor-dependent metabolic reprogramming of tolerogenic DCs is often closely linked to cellular stress responses^25^ ^26^. This was observed in our prior work demonstrating mregDC development to be driven by the sterol regulatory element-binding protein 2 (SREBP2) master regulator of the mevalonate biosynthetic pathway, a transcriptional program activated in response to lactate-induced pH stress within the tumor microenvironment (TME)^24^. Cellular adaptation to endoplasmic reticulum (ER) stress is achieved through activation of the unfolded protein response (UPR). The UPR is an integrated signal transduction pathway that conveys information about the folding state of ER proteins to the nucleus and cytoplasm^27^. Inositol-requiring enzyme 1 α (IRE1α) is the most conserved UPR stress sensor and acts as an endoribonuclease to process the mRNA of the transcription factor X-box binding protein-1 (XBP1), thereby generating the active spliced XBP1 isoform (XBP1s)^28^. Protein Kinase R-like Endoplasmic Reticulum Kinase (PERK) represents an alternative stress sensor that induces the transcriptional upregulation of activating transcription factor 4 (ATF4), another important regulator of the UPR^29^. Emerging evidence suggests that these pathways play a key role in regulating lipid metabolism and immune cell function in cancer^30–33^.

Many of the mediators that have been described to promote DC dysfunction in the TME exhibit short-range signaling activity and are unlikely to impact more distant DC populations. However, the DCs located within distant tissues like the TDLN play important roles in orchestrating anti-tumor immunity^34^ ^35^. LN tissues represent the primary anatomic site of DC-dependent antigen-presentation where they elicit adaptive immune responses through the activation of naïve CD4 ^+^ and CD8^+^ T cells^36^. Unlike many paracrine signaling mediators such as Wnt ligands and local metabolites, tumor extracellular vesicles (EVs) have the capacity to impact cell populations at sites distant from the tumor^37^. EVs are naturally released lipid-bilayer nanovesicles with a diameter between 30-150 nm^38^ ^39^. Previous studies have shown that tumor-derived EVs can promote tumor progression by delivering cargos containing genetic and molecular cargo to neighboring cells in the TME, as well as to distant cells in metastatic sites^40–42^. These studies have further implied that tumor-derived EVs mediate immune evasion, however, the underlying mechanisms have not been fully elucidated^43^.

Herein, we demonstrate that tumor-derived EVs compromise anti-tumor immunity by acting as long-range mediators and triggering a robust UPR in DCs that includes the activation of the PERK-ATF4 and IRE1α-XBP1 signaling axes. These pathways trigger SREBP2-dependent cholesterol biosynthesis and activation of the metabolic regulator peroxisome proliferator-activated receptor alpha (PPAR-α) to alter lipid homeostasis and drive FAO, respectively. Ultimately, the induction of these pathways by tumor-derived EVs promote a pro-tolerogenic DC phenotype that suppresses CD8^+^ T cell activation within the TME. Targeting this tumor EV-driven pathway offers a new therapeutic strategy to potentially overcome resistance to our current arsenal of checkpoint inhibitors.

## MATERIALS AND METHODS

### Experimental Mice

Mice were housed in pathogen-free facilities maintained by either the Duke University Division of Laboratory Animal Resources (DLAR) or the University of North Carolina Animal Care and Use Program. All experiments were performed in accordance with protocols approved by the Duke University Institutional Animal Care and Use Committee (IACUC) at Duke University and University of North Carolina, respectively. The following strains were purchased from The Jackson Laboratory and maintained in our animal colony: C57BL/6 (Strain #000664), OT-I (C57Bl/6-Tg (TcraTcrb)1100Mjb/J, Strain #003831), Foxp3-GFP (B6.Cg-Foxp3^tm2Tch^/J, Strain #006772), and BALB/c (Strain #000651). Additionally, for autochthonous melanoma studies, the inducible transgenic B6.Cg-Braf^tm1Mmcm^Pten^tm1Hwu^ Tg(Tyr-cre/ER^T^^2^)13Bos/BosJ mode (Common Name: BRAF^CA^;PTEN^loxP^;Tyr::CreER^T^^2^) was utilized to recapitulate the tumor immune microenvironment (TIME) and tumor-immune co-evolution of human melanoma. In this model, 4-hydroxytamoxifen (4-HT) was administered to induce *Braf^V600E^* expression and genetically ablate *Pten* in melanocytes. For syngeneic transplant experiments, a BRAF^V600E^PTEN^-/-^ melanoma cell line derived from resected melanoma tissues of the corresponding autochthonous model was implanted into the base of the tail of syngeneic C57BL/6 mice^44^. Zbtb46-Cre mice (B6.Cg-Zbtb46tm3.1(cre)Mnz/J, Strain #028538, The Jackson Laboratory) were crossed with Pparα^fl/fl^ mice (kindly provided by Dr. Frank A. Gomez, National Cancer Institute) to generate DC-specific *Ppar*α-deficient mice^45^. Wild-type (WT) or non-Cre-expressing littermates were utilized as controls.

### Tissue Culture

The BRAF^V600E^ PTEN^-/-^ melanoma cell line was generated by spontaneous immortalization of a tumor resected from the BRAF^CA^;PTEN^loxP^;Tyr::CreER^T^^2^ mouse model and cultured in DMEM supplemented with 10% FBS and 1% Antibiotic-Antimycotic (ThermoFisher Cat. #15240062)^44^. EO771 breast cancer cells were kindly provided by the laboratory of Dr. Donald P. McDonnell (Duke University). NIH/3T3 fibroblast cells were purchased from the UNC Lineberger Comprehensive Cancer Center Tissue Culture Facility. Both EO771 and NIH/3T3 cells were maintained in DMEM supplemented with 10% FBS and 1% Antibiotic-Antimycotic. The genetically engineered AKPS (APC^-/-^KRAS^G12D^p53^-/-^SMAD4^-/-^ colon cancer) cell line was kindly provided by the laboratory of Dr. Jatin Roper (Duke University) and was cultured in advanced DMEM (ADMEM) with 10% FBS 1× glutamine and 1× B27.

Bone marrow-derived dendritic cells (BMDCs) were generated from C57BL/6 or BALB/c mice. Bone marrow suspensions were harvested by flushing the tibias and fibulas with RPMI, followed by density gradient centrifugation using Lympholyte Cell Separation Media (Cedarlane Labs, CL5030) to remove stroma, erythrocytes, and debris. Cells were differentiated in RPMI supplemented with 10% FBS, 1% Antibiotic-Antimycotic, 2-mercaptoethanol (Gibco, 21-985-023), 20 ng/mL GM-CSF (Thermo Fisher Scientific, 315-03), and 10 ng/mL IL-4 (Thermo Fisher Scientific, 214-14). The culture medium was replenished every 48 hours (hrs). On day 7, BMDCs were harvested and purified via positive magnetic selection using CD11c MicroBeads (Miltenyi Biotec, Cat# 130-125-835).

### Extracellular Vesicle (EV) Isolation

BRAF^V600E^PTEN^-/-^ melanoma cells were expanded in T180 flasks in DMEM supplemented with 10% fetal bovine serum (FBS; Genesee Scientific, Cat# 25-550). Upon reaching 80% confluency, cells were passage into four 525 cm^2^ 3-layer Falcon® Multi-Flasks (VWR, Cat#353143) and cultured for 48 hrs. To eliminate contamination from bovine serum-derived EVs, the cell monolayers were washed three times with sterile PBS, and the medium was replaced with serum-free DMEM. Conditioned media was harvested after 48 hrs of serum-free culture. The collected supernatant was centrifuged at 2,000 x g for 20 minutes (mins) to pellet cells and large debris. The supernatant was then transferred to Beckman centrifuge tubes (Beckman Counter, Cat#355655) and spun at 10,000 x g on Beckman L70 for 30 mins to remove apoptotic bodies and microvesicles. The clarified supernatant was transferred to new tubes and ultracentrifuged at 100,000 x g for 90 mins using a Beckman Type 45 Ti fixed-angle rotor (Beckman Counter, Cat#339160) to pellet small EVs. The crude EV pellet was resuspended in sterile PBS and subjected to a second ultracentrifugation wash step at 100,000 x g for 90 mins to remove soluble protein contaminants. The final EV pellet was resuspended in 300 µl of sterile PBS^46^. EVs from EO771, AKPS and NIH/3T3 cell lines were isolated from conditioned media using the identical differential ultracentrifugation protocol described above. Protein concentration was quantified using a BCA Protein Assay Kit (Pierce, Thermo Scientific, Cat#PI23228).

### Extracellular Vesicle (EV) Characterization

To assess vesicle morphology and size, isolated EV suspensions were fixed in 2% paraformaldehyde (PFA). 5 µl of the fixed sample was deposited onto glow-discharged Formvar/carbon-coated copper grids (Electron Microscopy Sciences) and allowed to adsorb for 20 mins. Grids were washed with deionized water and negatively stained with 2% uranyl acetate for 1 min. Excess stain was removed and grids were air-dried. Imaging was performed using a FEI Tecnai G² Twin transmission electron microscope (Duke University Center for Electron Microscopy and Nanoscale Technology). Images were acquired at a range of magnifications up to 195,000x and expected EV morphology was verified. For size distribution analysis, the diameters of individual vesicles (n = 342, derived from random fields) were measured using ImageJ software to confirm the vesicular size range. Vesicle size data were binned in 10 nm increments and plotted as a frequency distribution histogram. The analysis revealed a heterogeneous population of EVs with a calculated mean diameter of 58.8 nm and a median diameter of 53.3 nm.

To validate EV purity, protein lysates were prepared from whole cells and isolated EVs using RIPA lysis buffer (Sigma-Aldrich, Cat#R0278) supplemented with protease and phosphatase inhibitors (Thermo Fisher, Cat#78441). Protein concentration was quantified via BCA assay. Equal amounts of protein (20µg) were resolved by SDS-PAGE and transferred onto PVDF membranes (Bio-Rad, Cat#1620177). Membranes were probed with primary antibodies overnight against positive exosomal markers, CD63 (Novus Biologicals, Cat#NBP2-34689; 1:1000), and CD81 (Santa Cruz, Cat#sc-23962; 1:200), and the negative control Golgi marker GM130 (to exclude cellular contamination; BD Transduction Laboratories, Cat#610822; 1:1000). Following incubation with HRP-conjugated anti-rabbit (Cell Signaling, Cat#70745) or anti-mouse (Sigma, Cat#GENXA931) secondary antibodies at 1:3000 for 1h at room temperature (RT), bands were visualized using an enhanced chemiluminescence (ECL) detection system Azure 300 (Azure Biosystems, Cat#AZI300) or Bio-Rad ChemiDoc Imaging System (Bio-Rad, Cat#12003153).

### Extracelluar Vesicle Quantification

EV abundance was quantified by measuring the activity of the exosome-enriched enzyme acetyl-CoA acetylcholinesterase (AChE) using the EXOCET Exosome Quantitation Kit (System Biosciences, Cat# EXOCET96A-1), following the manufacturer’s instructions. Briefly, isolated EV pellets were resuspended in the provided Lysis Buffer and incubated for 5 min on ice to release intraluminal AChE. Lysates and standards were loaded into a 96-well plate, followed by the addition of the Assay Reaction Buffer. The plate was incubated at room temperature for 25 min, and absorbance was measured at 405 nm using a microplate reader. The concentration of exosomes in each sample was calculated by interpolation from a standard curve generated using the provided exosome standards.

Based on the provided standard curve in **Supplementary Table S1**, the number of EVs per µg protein is 4.98 x 10^7^. A concentration of 20 µg/mL was selected to mimic physiological conditions, consistent with circulating exosome burdens reported in cancer patients (1∼ 6 x 10^9^ particles/ml)^47^. Based on the specific particle-to-protein ratio determined by EXOCET 4.98 x 10^7^ exosomes/µg), this dosage falls within the derived physiological range of approximately 10–60 µg/ml.

### Extracellular Vesicle (EV) Labeling and Tracking

To monitor EV uptake and biodistribution, purified BRAF^V600E^PTEN^-/-^ melanoma-derived EVs were labeled using the ExoGlow™-Protein EV Labeling Kit (Green) (System Biosciences, Cat# EQ800A-1). 300 µg of EV protein was resuspended in PBS and incubated with the labeling reaction mix for 20 mins at 37°C with gentle agitation. Unincorporated dye was removed using the provided Exosome Spin Columns (MWCO 3000) to ensure specific labeling of vesicular cargo. The resulting labeled EVs (designated "Exogreen^+^ EVs") were immediately used for downstream experiments.

### Tissue Digestion and Single-Cell Preparation

Tumors and lymph nodes (LNs) were resected, mechanically dissociated using a gentleMACS Dissociator (Miltenyi Biotec), and digested in RPMI containing 1 mg/ml Collagenase Type IV (Sigma-Aldrich, Cat# C5138), 20 U/ml DNase Type IV (Sigma-Aldrich, Cat# D5025) and 0.1 mg/ml Hyaluranidase (Sigma-Aldrich, Cat# H6254; tumor tissue only) for 20 mins at 37°C. Digested tissues were passed through 40 µM cell strainers and the reaction was quenched with RPMI supplemented with 10% FBS.

### Purification of DCs for *In Vitro* Assays

For *in vitro* immune cell assays, mouse spleens were freshly isolated, minced, and subjected to enzymatic dissociation in RPMI containing 1 mg/ml Collagenase Type IV (Sigma-Aldrich, Cat# C5138) and 20 U/ml DNase Type IV (Sigma-Aldrich, Cat# D5025). Single cell suspension were prepared by passing the digested tissue through a 40 µM cell strainer (VWR, Cat# 76327-098). The cells were then treated with RBC Lysis Buffer (Biolegend, Cat# 420301) for 5 min at RT and washed with PBS. Dendritic cells were subsequently isolated via positive magnetic selection using CD11c MicroBeads (Miltenyi Biotec, Cat# 130-125-835) according to the manufacturer’s protocol.

### Flow Cytometry

Single-cell suspensions were stained with Live/Dead Fixable Aqua or Violet Dead Cell Stain Kit (Thermo Fisher Scientific, Cat# L34957/L34955) at a 1:1000 dilution (1 ul/ml) in PBS for 30 mins at RT. Cells were washed twice in FACS buffer (PBS containing 5% FBS and 2 mM EDTA) and resuspended in anti-CD16/CD32 (Fc Block, BD Biosciences, Cat# 553142; 10 µg/mL) for 15 mins at 4°C. Fluorophore-conjugated primary antibodies were added directly to the blocking solution and incubated for 1 hour at 4°C. Cells were washed twice in FACS buffer, fixed using BD Cytofix (BD Biosciences, Cat# 554655) for 15 mins at 4°C, and washed two additional times. Data were acquired on a BD Canto/LSRFortessa X-20 flow cytometer at the Duke University Flow Cytometry Core Facility, an Attune NxT/LSRFortessa at the UNC Flow Cytometry Core Facility, or a Cytek Aurora spectral flow cytometer at UNC Lineberger Comprehensive Cancer Center.

Single-cell suspensions were seeded into a 96-well V-bottom plate. Cells were stained with Live/Dead Fixable Aqua or Violet Dead Cell Stain Kit (Thermo Fisher Scientific, Cat# L34957/L34955) at a 1:1000 dilution in PBS for 20 mins at RT. Cells were washed twice in FACS buffer and resuspended in anti-CD16/CD32 (Fc Block, BD Biosciences, Cat# 553142; 10 ug/ml) for 10 mins at 4°C. Fluorophore-conjugated primary antibodies (**Supplementary Table S2**) against surface markers were added directly to the blocking solution and incubated for 1 hour at 4°C. Cells were washed twice in FACS buffer. For experiments without intracellular staining, cells were washed twice in FACS buffer, fixed using BD Cytofix (BD Biosciences, Cat# 554655) for 15 mins at 4°C, and washed two additional times. For experiments with BODIPY/FAOBlue probes or GFP, samples were analyzed the same day. For intracellular staining, cells were then fixed and permeabilized using the eBioscience Foxp3/Transcription Factor Staining Buffer Set (Thermo Fisher Scientific, Cat#00-5523-00) according to the manufacturer’s instructions. Briefly, cells were incubated in Fixation/Permeabilization working solution for 1 hour at 4°C in the dark, washed twice with 1X Permeabilization Buffer, and stained with PE anti-mouse Foxp3 antibody in 1X Permeabilization Buffer overnight at 4°C. After staining, cells were washed twice with 1X Permeabilization Buffer and resuspended in FACS buffer. Data were acquired on a BD Canto or a BD LSRFortessa Flow Cytometer (Duke University Flow Cytometry Core Facility) or a Cytek Aurora Spectral Flow Cytometer (Lineberger Comprehensive Cancer Center, University of North Carolina).

### Annexin V Apoptosis Assay

To assess the cytotoxic effect of extracellular vesicles (EVs) on DCs, apoptosis rates were quantified using the eBioscience™ Annexin V Apoptosis Detection Kit (eFluor 450, 7-AAD; Thermo Fisher Scientific, Cat# 88-8006-72) according to the manufacturer’s instructions. Briefly, splenic DCs were seeded in a 96-well plate and incubated with 20 µg/ml of EVs or a PBS control for 24 hrs. Following treatment, cells were harvested, washed with cold PBS, and resuspended in 1X Binding Buffer. Cells were stained with fluorochrome-conjugated Annexin V and 7-AAD viability dye for 15 mins at RT in the dark. Samples were immediately acquired using BD FACSCANTO II Flow Cytometer (Duke University Flow Cytometry Core Facility). Data were analyzed using FlowJo software. Early apoptotic cells were defined as Annexin V^+^/7-AAD^-^, while late apoptotic cells were defined as Annexin V^+^/7-AAD^+^.

### Western Blot

Whole cell lysates (WCL) and exosome lysates were prepared using RIPA Lysis and Extraction Buffer (Thermo Fisher Scientific, Cat# 89901) supplemented with protease and phosphatase inhibitors (Roche, Cat# 4693159001, #4906845001). For nuclear protein analysis, BMDCs were treated with tumor-derived EVs (20 µg/ml) for 24 hrs. For PPAR nuclear translocation dose-response experiments, BMDCs were treated with varying concentrations of EVs (0.5, 2 or 10 µg/ml) for 24 hrs. For PERK inhibition experiments, BMDCs were pre-incubated with the PERK inhibitor GSK2606414 (500 µM; MedChemExpress, Cat# HY-18072) for 90 min prior to EV stimulation (20 µg/ml, 24 hrs).

Nuclear proteins were fractionated from BMDCs using the Nuclear Extraction Kit (Abcam, Cat# ab113474) according to the manufacturer’s protocol for suspension cells. Briefly, cells were harvested, washed with PBS, resuspended in 1X Pre-Extraction Buffer, and incubated on ice for 10 mins. After vigorous vortexing for 10 seconds, the suspension was centrifuged at 12,000 rpm for 1 min to separate the cytoplasmic fraction. The remaining nuclear pellet was resuspended in Extraction Buffer supplemented with DTT and Protease Inhibitor Cocktail (1:1000). Nuclei were lysed by incubation on ice for 15 mins with intermittent vortexing (5 seconds every 3 min), followed by sonication in an ice-cold water bath for three 10-second cycles. The lysate was centrifuged at 14,000 rpm for 10 min at 4°C, and the nuclear supernatant was collected. For analysis of IRE1α phosphorylation, DCs were serum-starved overnight and then treated with tumor-derived EVs (20 µg/ml) for 4 hrs. Tunicamycin (1µg/ml; Sigma-Aldrich, Cat# T7765) was used as a positive control. For IRE1α inhibition experiments, cells were pre-treated with the IRE1α inhibitor 4µ8c (10µM; EMD Millipore, Cat# 412512) for 20 min prior to the addition of EVs (20 µg/ml) for 24 hrs.

For immunoblotting, protein samples (20–30 µg for WCL, 30 µg for exosomes, or 10 µg for nuclear lysates) were resolved by SDS-PAGE and transferred to PVDF membranes (Bio-Rad, Cat# 1620177). Membranes were blocked and incubated with primary antibodies, followed by horseradish peroxidase (HRP)-conjugated secondary antibodies (diluted 1:3000 in 5% non-fat milk for 1 hour at RT). Primary antibodies included: Rab27a (Cell Signaling Technology [CST], Cat# 69295S), SREBP2 (Novus Biologicals, Cat# NB100-74543), ATF4 (CST, Cat# 11815S), ATF6 (CST, Cat# 65880S), IRE1α (CST, Cat# 3294S), SREBP1 (clone 2A4; Santa Cruz Biotechnology, Cat# sc-13551), Lamin B1 (clone B-10; Santa Cruz, Cat# sc-374015), XBP-1s (CST, Cat# 40435), PPARα (Thermo Fisher Scientific, Cat# PA1-822A), PPARγ (Santa Cruz, Cat# sc-7273), PPARβ (Santa Cruz, Cat# sc-74517) and Histone H3 (Santa Cruz, Cat# sc-517576). Unless otherwise indicated, primary antibodies were diluted 1:1000 in 5% non-fat dry milk (Bio-Rad, Cat# 1706404) and incubated overnight at 4°C. Exceptions included phospho-IRE1α (Ser724) (Novus Biologicals, Cat# 2323s), which was diluted 1:100 in 1% BSA, PPARγ (1:200), PPARβ (1:100), Lamin B1 (1:100) and HRP-conjugated β-Actin (clone C4; Santa Cruz, Cat# sc-47778), which was incubated for 1 hour at RT. Immunoreactive bands were visualized using SuperSignal™ West Dura or Femto Extended Duration Substrate (Thermo Scientific, Cat# 34076 or #34905) and imaged using the ChemiDoc MP Imaging System (Bio-Rad, Cat# 170-8280) or the Azure Imaging System (Azure Biosystems, Cat# AZI300).

### Generation of Rab27a Knockdown (KD) Cell Lines

Bacterial glycerol stocks of shRNA constructs targeting mouse Rab27a (TRCN0000100575, TRCN0000100577, and TRCN0000100578) and a non-targeting scramble control (SHC002) were obtained from the Sigma-Aldrich MISSION® shRNA library. To isolate individual clones, bacterial stocks were enriched in SOC media (Thermo Fisher, Cat# 15544034) and streaked onto Luria-Bertani (LB) agar plates containing 100 µg/ml carbenicillin. Single colonies were expanded, and plasmid DNA was isolated using a Miniprep Kit (Qiagen, Cat#27104). Plasmid sequence identity was confirmed via whole-plasmid sequencing (Plasmidsaurus). Lentiviral particles were produced in HEK293T cells by co-transfection of the purified pLKO.1-puro transfer plasmid with the packaging plasmid psPAX2 and the envelope plasmid pMD2.G. Transfection was performed using Lipofectamine 3000 (Thermo Fisher, Cat# L3000015). Viral supernatants were harvested 48 hours post-transfection, centrifuged to remove cell debris, and filtered through a 0.45 µm PES filter. Target tumor cells were transduced with the collected lentiviral supernatants in the presence of 8 µg/ml polybrene (hexadimethrine bromide) at approximately 50% confluency to maximize killing efficiency during selection. Transduced cells were selected using 2.5 ug/ml puromycin for 3–4 days, followed by a maintenance dose of 1 µg/ml. Knockdown efficiency was validated by Western blot analysis.

### Quantitative Real-Time Polymerase Chain Reaction (qrt-PCR)

Total RNA was isolated from DCs using either TRIzol Reagent (Thermo Fisher Scientific, Cat# 15596026) or the RNeasy Plus Mini/Micro kits (Qiagen, Cat# 74134, #74004), following the manufacturers’ instructions. cDNA was synthesized using the iScript cDNA Synthesis Kit (Bio-Rad, Cat# 1708890). For low-input reactions, cDNA templates were pre-amplified using the SsoAdvanced PreAmp Supermix (Bio-Rad, Cat# 1725160). qRT-PCR was performed using Power SYBR Green PCR Master Mix (Thermo Fisher Scientific, Cat# 4367659) on an Applied Biosystems 7500 Real-Time PCR instrument. Relative gene expression was analyzed using the comparative C_t_ (2^−ΔΔ𝐶𝑡^) method. The specific sequences for all qPCR and genotyping primers used in this study are provided in **Supplementary Table S3**.

### RNA Sequencing and Gene Expression Analysis

For transcriptional profiling, RNA was isolated from cultured BMDCs using the RNeasy Mini Kit (Qiagen) according to the manufacturer’s instructions. For FACS-isolated lymph node-derived DCs (LNDCs), RNA was extracted using TRIzol™ Reagent (Thermofisher, Cat# 15596026), followed by purification using the RNeasy Micro Kit (Qiagen). RNA samples were sent to Novogene (Novogene Corporation Inc., Sacramento, CA, USA) for library preparation and sequencing. For BMDC samples, messenger RNA (mRNA) libraries were prepared with poly(A) enrichment and sequenced on an Illumina NovaSeq X Plus platform (paired-end, 150 bp). For sorted ExoGreen^+^ LNDC samples, RNA was amplified using the SMARTer® Kit followed by ultra-low input mRNA non-directional library preparation. These libraries were sequenced on an Illumina NovaSeq X Plus platform (paired-end, 150 bp). Raw sequencing reads were quality-checked and trimmed for adapters and low-quality bases. The processed reads were aligned to the mouse reference genome (mm10) using the STAR aligner. Gene expression quantification and differential expression analysis were performed using the DESeq2 R package. Genes with an adjusted p-value < 0.05 and a fold change > 1.5 were considered significantly differentially expressed.

### scRNA-seq Library Preparation and Sequencing

DCs were isolated from spleens using CD11c MicroBeads (Miltenyi Biotec) as described above. Single-cell suspensions were encapsulated into Gel Bead-in-Emulsions (GEMs) using the Chromium Next GEM Single Cell 3’ Kit v3.1 (10x Genomics, Cat# PN-1000269) according to the manufacturer’s instructions. Library quality was assessed using an Agilent 2100 Bioanalyzer, and final libraries were quantified using the NEBNext Library Quant Kit for Illumina (New England Biolabs, Cat# E7630L). Sequencing was performed at the Duke University Sequencing and Genomic Technology Core Facility on an Illumina NovaSeq 6000 at a target depth of ∼40,000 read pairs per cell.

### PPARα Transcription Factor Activity Assay

BMDCs were generated as described above and seeded at a density of 1 x 10^6^ cells/ml in 48-well plates. Cells were pre-treated with the IRE1α inhibitor 4u8c (10 µM; stock concentration 100 mM) or vehicle control (DMSO) for 20 mins, followed by stimulation with 20 ug/ml melanoma EVs for 24 hrs. Nuclear extracts were subsequently prepared using the Nuclear Extraction reagents included in the PPAR-α Transcription Factor Assay Kit (Abcam, Cat # ab133107). PPAR-α DNA-binding activity was quantified by adding equal amounts of nuclear extract to a 96-well plate immobilized with an oligonucleotide containing the peroxisome proliferator response element (PPRE). The assay was performed strictly according to the manufacturer’s instructions. Following incubation with the PPAR-α primary antibody and HRP-conjugated secondary antibody, the developing solution was added, and the reaction was stopped. Absorbance was measured at 450 nm using a microplate reader. Relative PPAR-α activity was calculated by normalizing the OD450 values to the untreated control group.

### Fatty Acid Oxidation Assay

To assess fatty acid oxidation (FAO) activity, splenic DCs were isolated as described above and seeded at a density of 2 x 10^5^ cells/200 µl in 96-well plates. Cells were treated with 20 µg/ml melanoma EVs and incubated overnight. Following surface antibody staining, cells were washed and resuspended in HEPES buffered saline (Sigma-Aldrich, Cat #H0887) containing the FAOBlue detection reagent (DiagnoCine, Cat# FNK-FDV-0033) diluted 1:2000 (5 µM final concentration) is a non-fluorescent probe that is oxidized by the FAO pathway into a fluorescent product, with the resulting mean fluorescence intensity (MFI) being proportional to FAO activity. Cells were incubated for 30 mins at 37°C. After washing, cells were immediately analyzed on a BD LSRFortessa flow cytometer (Duke University Flow Cytometry Core Facility). FAO activity was quantified by calculating the mean fluorescence intensity (MFI) of the FAOBlue signal in the Pacific Blue channel.

### SCENITH Metabolic Profiling Assay

TDLNs and tumors were harvested and dissociated into single-cell suspensions as described above. Cells were washed and plated at a density of 1 x 10^6^ cells/ml in RPMI 1640 supplemented with 1 mM pyruvate, 2 mM glutamine, and 10 mM glucose. To profile metabolic activity, cells were treated with vehicle control or the following metabolic inhibitors for 45 minutes at 37°C: 2-Deoxy-D-glucose (2-DG; 100 mM) to inhibit glycolysis, Oligomycin (1 µM) to inhibit mitochondrial respiration, or Etomoxir (40 µM) to inhibit fatty acid oxidation. Puromycin (10 µg/ml; Sigma-Aldrich) was added to the culture for the final 15 minutes of the incubation period. Cells were washed with cold PBS and stained with Live/Dead Fixable Aqua Dead Cell Stain Kit (Thermo Fisher Scientific). Following Fc blockade, cells were stained for surface markers to identify dendritic cell populations. Cells were then fixed and permeabilized using the eBioscience Foxp3/Transcription Factor Staining Buffer Set (Thermo Fisher Scientific). Intracellular staining was performed for 1 hour using an anti-Puromycin antibody (Clone 3RH11, Kerafast, Cat# Kf-Ab02366-1.1 (EQ0001)) that was labeled using the Alexa Fluor® 594 Conjugation Kit (Fast) - Lightning-Link® (Abcam, Cat# ab269822). Briefly, the antibody was mixed with the Modifier reagent and incubated for 15 minutes, followed by the addition of the Quencher reagent for 5 minutes prior to use. Data were acquired on a BD LSRFortessa (Duke University Flow Cytometry Core Facility). Metabolic capacities (Mitochondrial and FAO) were calculated based on the percentage of protein synthesis inhibition (reduction in Puromycin MFI) in inhibitor-treated cells relative to the vehicle control^48^.

### BODIPY Lipid Content Analysis

To assess neutral lipid content and lipid peroxidation, splenic DCs were seeded at a density of 2 x 10^5^ cells/200 µl in 96-well plates. Cells were treated with tumor-derived EVs (20 µg/ml) in the presence or absence of metabolic inhibitors, including the SREBP2 inhibitor Fatostatin (25µM), the PPAR-α antagonist TPST-1120 (20µM), the PPARδ antagonist GSK 3787 (10 µM), or the PPARγ antagonist GW 9662 (10 µM) and incubated overnight. As a positive control for lipid peroxidation, a separate group of cells was treated with 300µM hydrogen peroxide (H_2_O_2_) for 30 minutes prior to staining. Following treatment, cells were washed and resuspended in PBS containing either BODIPY 493/503 (Thermo Fisher Scientific, Cat# D3922) or BODIPY 581/591 C11 (Thermo Fisher Scientific, Cat# D3861) at a final concentration of 500 ng/ml. Cells were incubated for 30 minutes at 37°C, washed twice in PBS, and resuspended in FACS buffer for immediate acquisition on a BD LSRFortessa (Duke University Flow Cytometry Core Facility). Neutral lipids (BODIPY 493/503) were detected using a 488 nm excitation laser with a 525/50 bandpass filter. For the analysis of lipid peroxidation (BODIPY 581/591 C11), the reduced form was measured using a 561 nm excitation laser with a 586/15 bandpass filter, while the oxidized form was measured using a 488 nm excitation laser with a 525/50 bandpass filter.

### TF2-C12 Fatty Acid Uptake Assay

Fatty acid uptake measurement in DCs were performed using a dodecanoic acid fluorescent TF2- C12 fatty acid (Sigma, Cat# MAK156) according to the manufacturer’s protocol. Briefly, BMDCs were treated with EVs (10 µg/ml) for 24h. Following treatment, cells were washed and resuspended in serum-free medium and serum-starved for 1 hour at 37°C to deplete endogenous lipids. Subsequently, 100 µg/ml of the prepared TF2-C12 Fatty Acid Dye Loading Solution was added to the cells and the plate was incubated for 60 minutes at 37°C protected from light. Fatty acid uptake was immediately analyzed by flow cytometry in the FITC channel (Ex=485nm/Em=515nm).

### *In Vitro* Treg Differentiation Assay

Splenic DCs were isolated from Balb/c mice (H-2^d^) as described previously. DCs were incubated in the presence or absence of tumor-derived EVs (20 µg/ml) overnight. Following incubation, DCs were washed twice with PBS and co-cultured with allogeneic naïve CD4^+^ T cells magnetically isolated from Foxp3-GFP reporter mice (H-2^b^) using the EasySep Mouse Naïve CD4^+^ T Cell Isolation Kit (Stem Cell Technologies, Cat# 19765). Co-cultures were maintained at a 1:5 DC:T cell ratio in RPMI-1640 supplemented with 10% FBS, 1% Antibiotic-Antimycotic (Sigma, Cat# A5955), 2-mercaptoethanol (Thermo Fisher Scientific, Cat# 21985023), and 200 pg/ml TGFβ (R&D Systems, Cat# 7666-MB-005/CF). Fresh medium was added on day 3, and on day 5, the frequency of Foxp3^+^ cells was assessed by same-day flow cytometry.

To further investigate the intrinsic role of PPAR-α in this process, a parallel mixed lymphocyte reaction (MLR) was performed using DCs isolated from Zbtb46-Cre x Pparα^fl/fl^ mice or their littermate controls (H-2^b^). These DCs were co-cultured with allogeneic naïve CD4^+^ T cells magnetically isolated from BALB/c mice and maintained under the identical polarizing conditions described above (1:5 ratio, 200 pg/ml TGFβ). On day 5, cells were harvested, fixed, and permeabilized using the eBioscience Foxp3/Transcription Factor Staining Buffer Set (Thermo Fisher Scientific, Cat#00-5523-00), and the frequency of Foxp3^+^ cells within the CD4^+^ T cell population was assessed by flow cytometry.

### *In Vivo* Treg Differentiation Assay

Splenic DCs were isolated from C57BL/6 mice (H-2^b^) and treated with tumor-derived EVs or PBS control overnight as described above. DCs were washed twice with PBS and resuspended at a concentration of 1 x 10^6^ cells/ml. A total of 5 x 10^4^ DCs in 50µl of PBS were injected intradermally into the footpads of Foxp3-GFP reporter mice (H-2^b^). Draining inguinal LNs were harvested 6–7 days post-injection. Single-cell suspensions were prepared and analyzed for the expansion of CD4^+^Foxp3^+^ regulatory T cells by same-day flow cytometry.

### OT-I T Cell Proliferation Assay

To determine the ability of EV-treated DCs to cross-present tumor-associated antigens, DCs were isolated and cultured in the presence or absence of tumor-derived EVs (20µg/ml) overnight. Subsequently, DCs were pulsed with ovalbumin (OVA) protein (Sigma-Aldrich) at a final concentration of 1 mg/ml for 6 hrs and washed extensively to remove free antigen. Naïve CD8^+^ T cells were isolated from OT-I mice (R&D Systems, Cat# MAGM207) and labeled with CellTrace CFSE (Thermo Fisher Scientific, Cat# C34554). For labeling, isolated T cells were washed once in serum-free RPMI (300 x g, 5 min). A staining solution was prepared by adding 9 µl of the reconstituted CFSE stock (dissolved in 18 µl DMSO) to 10 ml of serum-free RPMI. The cell pellet was rapidly resuspended in 10 ml of the staining solution and incubated for 20 minutes at 37°C protected from light. Labeling was quenched by washing the cells twice with RPMI supplemented with 10% FBS and β-mercaptoethanol (300 x g, 5 min). Labeled T cells were resuspended at a density of 5 x 10^5^ cells/ml, and 100 µl (50,000 cells) was plated per well in 96-well V-bottom plates. Co-cultures were initiated by adding 10,000 OVA-pulsed DCs (5:1 T cell:DC ratio) and maintained for 60-66 hrs. T cell proliferation and activation were assessed by flow cytometry analyzing CFSE dilution and surface CD44 expression, respectively. Notably, DCs were also pulsed with OVA protein prior to tumor EV exposure with no differences ultimately observed in CD8^+^ T cell proliferation responses.

### Mouse Tumor Studies

To determine the immune-dependence of EV-mediated tumor promotion, 1 x 10^6^ Rab27a-silenced BRAF^V600E^PTEN^-/-^ melanoma cells (BRAF^V600E^PTEN^-/-^-Rab27a^KD^) and non-target control BRAF^V600E^PTEN^-/-^ melanoma cells (BRAF^V600E^PTEN^-/-^-NTC) were implanted subcutaneously into immunocompetent C57Bl/6 mice and immunodeficient NOD.Cg-Prkdc^scid^ Il2rg^tm1Wjl^/SzJ (NSG) mice. Primary tumor volume measurements were performed for each group every 3 days by orthagonal caliper measurements. Tumor volume was determined based on the following formula: volume = [(length \times width^2^)]/2.

To assess the interaction between tumor-derived EVs and DC PPAR-α signaling, 1 x 10^6^ BRAF^V600E^PTEN^-/-^-Rab27a^KD^ and control BRAF^V600E^PTEN^-/-^-NTC melanoma cells were implanted at the base of the tail of DC-specific PPAR-α knockout mice (Ppara^fl/fl^Zbtb46-Cre) or their Zbtb46-Cre littermate controls. Primary tumor volume measurements were performed for each group every 3 days by orthagonal caliper measurements as above.

For authochtonous melanoma experiments, tumors were induced by injecting B6.Cg-Tg(Tyr-cre/ER^T^^2^)13Bos BRAF^tm1Mmcm^ Pten^tm1Hwu^/BosJ (BRAF^V600E^PTEN^-/-^ Tyr::CreER^T^^2^; Strain #013590) mice subdermally with 4-hydroxytamoxifen (4-HT; Millipore Sigma, H6278; 38.75 µg) at the base of the tail. When tumors reached an approximate volume of 75 mm³, mice were randomized into 4 treatment groups: 1) anti-IgG isotype control (200 μg) delivered by intraperitoneal (i.p.) injection every 3 days + vehicle control, 2) anti-PD-1 antibody (clone RMP1-14, 200 μg) i.p. every 3 days + vehicle control, 3) anti-IgG isotype control (200 μg) i.p. every 3 days + TPST-1120 (30 mg/kg in 0.5% methylcellulose) twice daily via orogastric (og) lavage, and4) anti-PD-1 antibody (200 μg) i.p. every 3 days + TPST-1120 (30 mg/kg) twice daily via og lavage.

Primary tumor volumes were monitored as described above. At the experimental endpoint, tumors and draining LN tissues were harvested and processed into single-cell suspensions for immunophenotyping. Flow cytometry was performed to analyze the infiltration and activation status of immune cell populations. Additionally, DCs were isolated from tumor and LN tissues to assess their metabolic profile using SCENITH analysis described above^48^. To quantify pulmonary metastasis, lungs were harvested at the experimental endpoint and metastatic burden was assessed by qRT-PCR analysis of the melanocyte differentiation antigen Trp2 (dopachrome tautomerase) as previously performed^49^. Similar studies were performed in syngeneic C57BL/6 hosts by implanting 1 x 10^6^ BRAF^V600E^PTEN^-/-^ melanoma cells were at the base of the tail.

### Computational Analysis

Raw sequencing data were demultiplexed and processed using the Cell Ranger 4.0.0 pipeline (10x Genomics). The cellranger mkfastq command was used to convert Illumina BCL files to FASTQ format. Subsequently, cellranger count was used to align reads to the mouse (mm10) (GRCh38; GENCODE v32/Ensembl 98) reference genomes, filter unique molecular identifiers (UMIs), and generate gene-barcode matrices.

Downstream analysis was performed in R using the Seurat package. Cells were filtered to exclude low-quality barcodes based on the following criteria: genes expressed in fewer than 3 cells, cells containing fewer than 200 or more than 5000 detected genes, and cells with >10% mitochondrial gene content. Post-filtering, data were log-normalized using the *NormalizeData* function and scaled using the *ScaleData* function to reduce the influence of highly expressed genes. To identify enriched pathways, Gene Set Enrichment Analysis (GSEA) was performed on differentially expressed genes using the *fgsea* R package. Enrichment was evaluated using gene sets from the Molecular Signatures Database (MSigDB), including the Hallmark collection and curated Reactome pathways (e.g., Regulation of Lipid Metabolism by PPARα), with statistical significance determined by adjusted p-values. The SREBP2 target gene signature used for heatmap visualization was derived from curated literature.

### Statistical Analysis

Graphpad Prism 10 (Graphpad Software) was used for all statistical analysis. An unpaired 2-tailed Student’s *t* test was used to compare mean differences between two groups. For data normalized to a control group (theoretical mean of 1.0), statistical significance was determined using a one-sample t-test. A one-tailed one-sample t-test was specifically utilized when testing a directional hypothesis (e.g., reduction of CD63). For comparisons involving three or more groups, a univariate one-way ANOVA followed by Sidak’s post hoc test (for planned comparisons) or a non-parametric Kruskal-Wallis test followed by Dunn’s multiple comparison test was performed. For tumor growth kinetics and grouped data sets involving two independent variables, a two-way ANOVA with multiple comparisons test was utilized. A *P* value of less than 0.05 was considered significant. Quantitative data was presented as mean ± SEM. Asterisks represent statistical significance (*P < 0.05, **P < 0.01, ***P < 0.001, ****P < 0.0001).

### Study Approvals

All mouse experiments were performed according to Institutional Animal Care and Use Committee (IACUC) approved protocols at Duke University Medical Center and the University of North Carolina at Chapel Hill Lineberger Comprehensive Cancer Center (LCCC).

## RESULTS

### Tumor-derived EVs Compromise DC Function and Impair Anti-Tumor Immunity

Previous studies have suggested that tumor-derived EVs contribute to the generation of an immunosuppressive TME^39–41^ ^50^ ^51^. Consistent with these findings, we also found that the genetic silencing of the Rab27a small GTPase known to be necessary for EV secretion, suppresses BRAF^V600E^PTEN^-/-^ melanoma progression only in immunocompetent mice (**figure 1A, online supplemental figures 1A-B**). To further characterize the immunologic impact of tumor EVs, using multi-parameter flow cytometry we also observed an increase in tumor-infiltrating CD8^+^ T cells and a concurrent reduction in CD4^+^FoxP3^+^ regulatory T cells (Tregs) in Rab27a-silenced melanomas (**figure 1B, online supplemental figure 1C**). This shift in the immune landscape suggested that tumor-derived EVs promote tumor progression at least partially by suppressing the development of anti-tumor immunity. Previous studies have established that myeloid lineages are the predominant recipients of tumor-derived EVs within the tumor microenvironment (TME)^52^. This suggests that the observed immune suppression is driven by the EV-mediated reprogramming of the myeloid populations within the TME. Given the critical role of DCs in orchestrating anti-tumor immunity as well as the ability of EVs to genetically modulate distant cell populations, we became interested in understanding the impact of tumor EVs on DC function and what role this process plays in driving tumor-mediated immune evasion.

**Figure 1.**
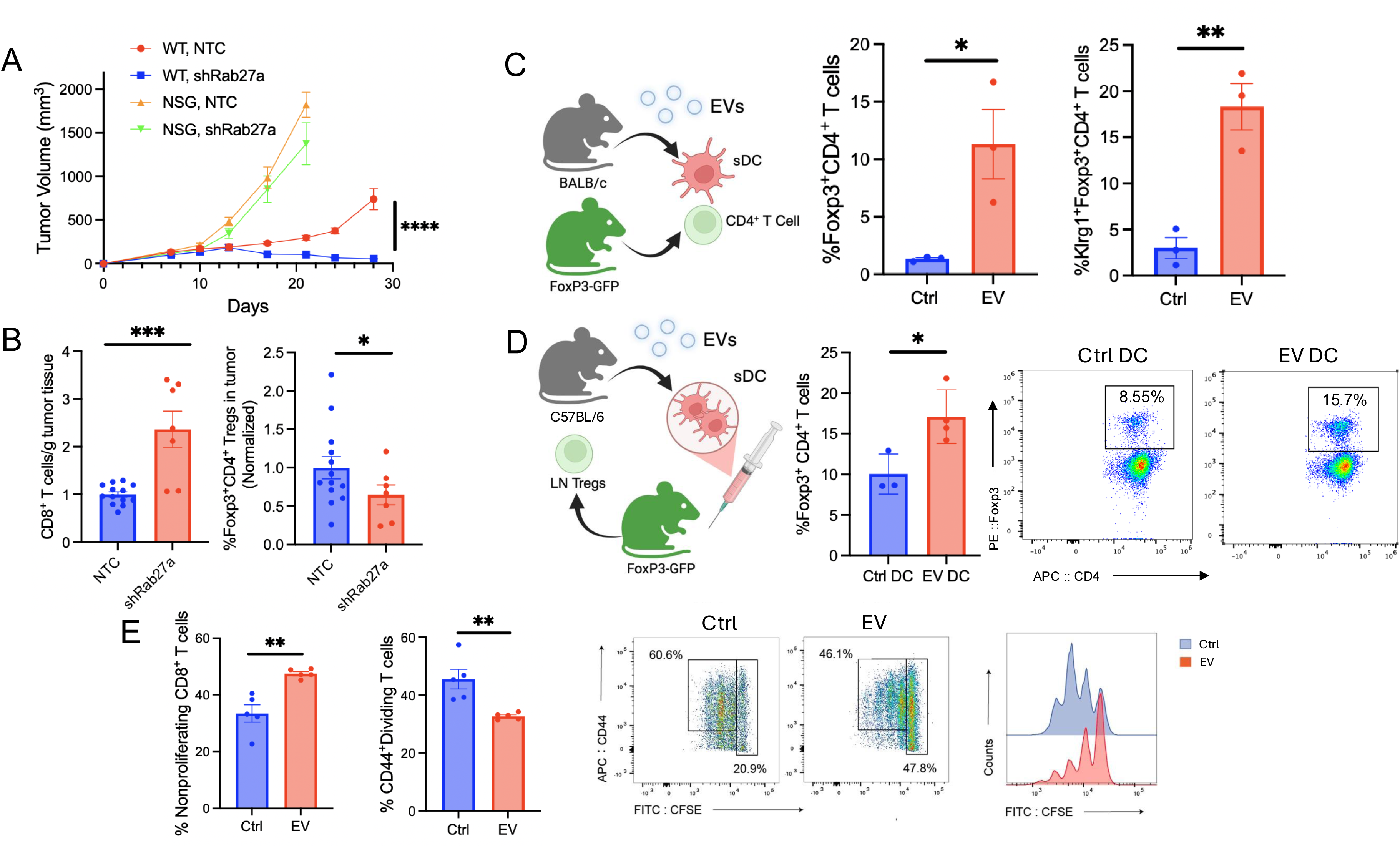
Tumor-derived EVs impair T cell function to promote tumor immune escape. **(A)** Tumor growth kinetics in syngeneic WT and NSG mice following implantation with BRAF^V600E^PTEN^-/-^ melanoma cells expressing non-targeting control shRNA (NTC) or Rab27a-targeted shRNA (shRab27a) over 28 days (n=8). Representative of two independent experiments. **(B)** Flow cytometry quantification of tumor-infiltrating lymphocytes in BRAF^V600E^PTEN^-/-^-NTC or BRAF^V600E^PTEN^-/-^-shRab27a melanomas in WT mice. CD8^+^ T cells normalized to tumor weight (*left*) and frequency of CD4^+^FoxP3^+^ Tregs (*right*) are shown as a percentage of total live CD45^+^ cells normalized to BRAF^V600E^PTEN^-/-^-NTC melanomas (n=7-13). Data aggregated from two independent experiments. **(C)** EV impact on *in vitro* DC-dependent Treg differentiation. Naive CD4^+^ T cells isolated from Foxp3-GFP reporter mice were co-cultured with allogeneic splenic DCs treated with PBS (Ctrl) or EVs (20 µg/ml) overnight. Shown are the quantification of *de novo* induced CD4^+^FoxP3^+^ Tregs (*left*) and KLRG1^+^CD4^+^FoxP3^+^ Tregs (*right*) (n=3). Representative of three independent experiments. **(D)** EV impact on *in vivo* DC-mediated Treg activation. BALB/c splenic DCs were treated with PBS (Ctrl) or EVs (20 µg/ml, 24 h) and injected into the footpads of syngeneic Foxp3-GFP reporter mice. *Left*, Draining LNs were harvested on days 6–7 for flow cytometry quantification of the frequency of KLRG1^+^ CD4^+^FoxP3^+^ Tregs in each group. *Right*, representative flow cytometry plots of KLRG1^+^CD4^+^FoxP3^+^ Tregs following delivery of DCs treated with PBS versus EVs (n=6-8). Representative of two independent experiments. **(E)** EV impact on DC-mediated CD8^+^ T cell activation. CFSE-labeled naïve CD8^+^ T cells were co-cultured with OVA-loaded splenic DCs following treatment with PBS (Ctrl) or EVs (20 µg/ml) overnight. Shown are the percent of non-proliferating (CFSE^high^) and proliferating (CFSE^low^CD44^+^) cells (*left/middle*) with representative flow cytometry plots and histograms (*right*) (n=5). Representative of three independent experiments. All data are reported as mean ± SEM. \**p*<0.05, \*\**p*<0.01, \*\*\**p*<0.001 by unpaired, two-tailed Student’s t-test or one-tailed one-sample *t*-test (B). See also online supplemental figure S1. **DC**, dendritic cell; **EV**, extracellular vesicle; **KLRG1**, killer cell lectin-like receptor G1; **LN**, lymph node; **NTC**, non-targeting control; **CFSE**, carboxyfluorescein succinimidyl ester; **shRNA**, short hairpin RNA; **Treg**, regulatory T cell; **WT**, wild-type, **NSG**, NOD *scid* gamma

Our prior work, including that of others, has broadly characterized tolerogenic DCs as exhibiting two defining functional hallmarks: 1) impaired antigen cross-presentation and 2) increased Treg expansion^22^ ^53^. To investigate whether tumor EV exposure induces a pro-tolerogenic DC phenotype, we initially isolated and characterized EVs from the BRAF^V600E^PTEN^-/-^ melanoma cell line using sequential ultracentrifugation, co-cultured EV-treated DCs with naïve CD4^+^ T cells, and monitored the differentiation of CD4^+^FoxP3^+^ Tregs *in vitro* by flow cytometry (**online supplemental figures 1D-F**). This analysis indicated tumor EV-treated DCs promote both the differentiation and activation of Tregs (**figure 1C, online supplemental figure 1G**)^54^ ^55^. This effect was also observed using EVs isolated from the EO771 breast cancer cell line, indicating that this phenomenon was not restricted to EVs isolated from melanoma cells (**online supplemental figure 1H**). In contrast, EVs isolated from non-tumorigenic NIH-3T3 fibroblasts did not promote DC-dependent Treg differentiation, suggesting that this effect may be restricted to malignant tissues (**online supplemental figure 1I)**. To further assess whether EV-dependent DC tolerization could also be observed *in vivo*, we delivered tumor EV-treated DCs into the footpad of FoxP3-GFP mice and found an increase in FoxP3-GFP^+^ Tregs in resected popliteal draining LN tissues (**figure 1D**). In parallel with Treg expansion, we also found tumor EV-treated DCs to exhibit an impaired ability to stimulate CD8^+^ T cell activation and proliferation, while no impact on DC viability was observed (**figure 1E, online supplemental figure 1J, K**). Similar to above, this suppressive phenomenon was recapitulated using EO771-derived EVs, while 3T3-derived EVs failed to impair CD8^+^ T cell proliferation (**online supplemental figures 1L, M**). Taken together, these observations suggest that tumor-derived EVs contribute to DC dysfunction, prompting us to further characterize the transcriptional program and phenotype of tumor EV-educated DCs.

### Tumor EVs Contribute to the Development of mregDCs

We have demonstrated that a population of DCs *in situ,* previously described as ‘mature DCs enriched in immunoregulatory molecules’ (mregDCs), exhibit a pro-tolerogenic function that supports melanoma progression^23^ ^24^. This work further revealed that the development of this mregDC population is partially dependent on the activation of sterol response element binding protein-2 (SREBP2) by tumor-derived lactate^24^. However, these observations indicated that additional TME-derived stimuli may also influence mregDC development and function. Consequently, we sought to determine if tumor EVs may contribute to mregDC development. To examine this question, we performed both bulk RNA sequencing (RNAseq) on bone marrow-derived DCs (BMDCs) and single-cell RNAseq (scRNAseq) analysis on splenic DCs treated with EVs isolated by sequential ultracentrifugation from the BRAF^V600E^PTEN^-/-^ melanoma cell line. Interestingly, both bulk RNAseq and scRNAseq analysis revealed tumor-derived EVs to induce a gene expression signature strikingly similar to the previously reported gene program of mregDCs (**figure 2A,B, online supplemental figure 2A**)^56^ ^57^. This was found to be consistent with our flow cytometry analysis showing tumor-derived EVs to induce the upregulation of several surface maturation markers by DCs (**online supplemental figure 2B**). Together, these data suggest that melanoma-derived EVs transcriptionally re-program DCs towards an immunoregulatory phenotype *in vitro*. To determine whether tumor-derived EVs may have a similar impact on DCs within the draining lymph node (LN) tissues *in vivo*, we delivered Exo-Green labeled tumor-derived EVs by intra-dermal injection into the footpad of syngeneic mice. Ultra-low input whole transcriptome RNAseq analysis was performed on Exo-Green^+^ and Exo-Green^-^ DCs isolated by FACS from draining LN tissues. In line with our previous findings, we observed a similar mregDC gene program enriched in Exo-Green^+^ DCs within the *in situ* LN microenvironment (**figure 2C, online supplemental figures 2C, D**). To further verify that tumor EVs can re-program tumor-associated DCs in a similar manner *in vivo*, we engineered a CD81-mEmerald fluorescent fusion protein-expressing BRAF^V600E^PTEN^-/-^ melanoma model. CD81 is a canonical tetraspanin marker that enriches on the surface of EVs, allowing for the monitoring of DC populations that internalize tumor-derived EVs. Indeed, we found CD81-mEmerald⁺ DC populations that have engulfed tumor EVs to exhibit a significant upregulation in the expression of key immunoregulatory genes associated with the mregDC signature (*Cd274*, *Pdcd1lg2*, *Cd200*, *Fas*, and *Socs1*) (**figure 2D, online supplemental figure 2E**), further suggesting that tumor-derived EVs induce a transcriptional program previously associated with mregDCs. In light of the pro-tolerogenic function of DCs exposed to tumor EVs discussed above, these cumulative findings suggest that tumor EVs serve as an additional stimulus capable of driving mregDC development in the TME.

**Figure 2.**
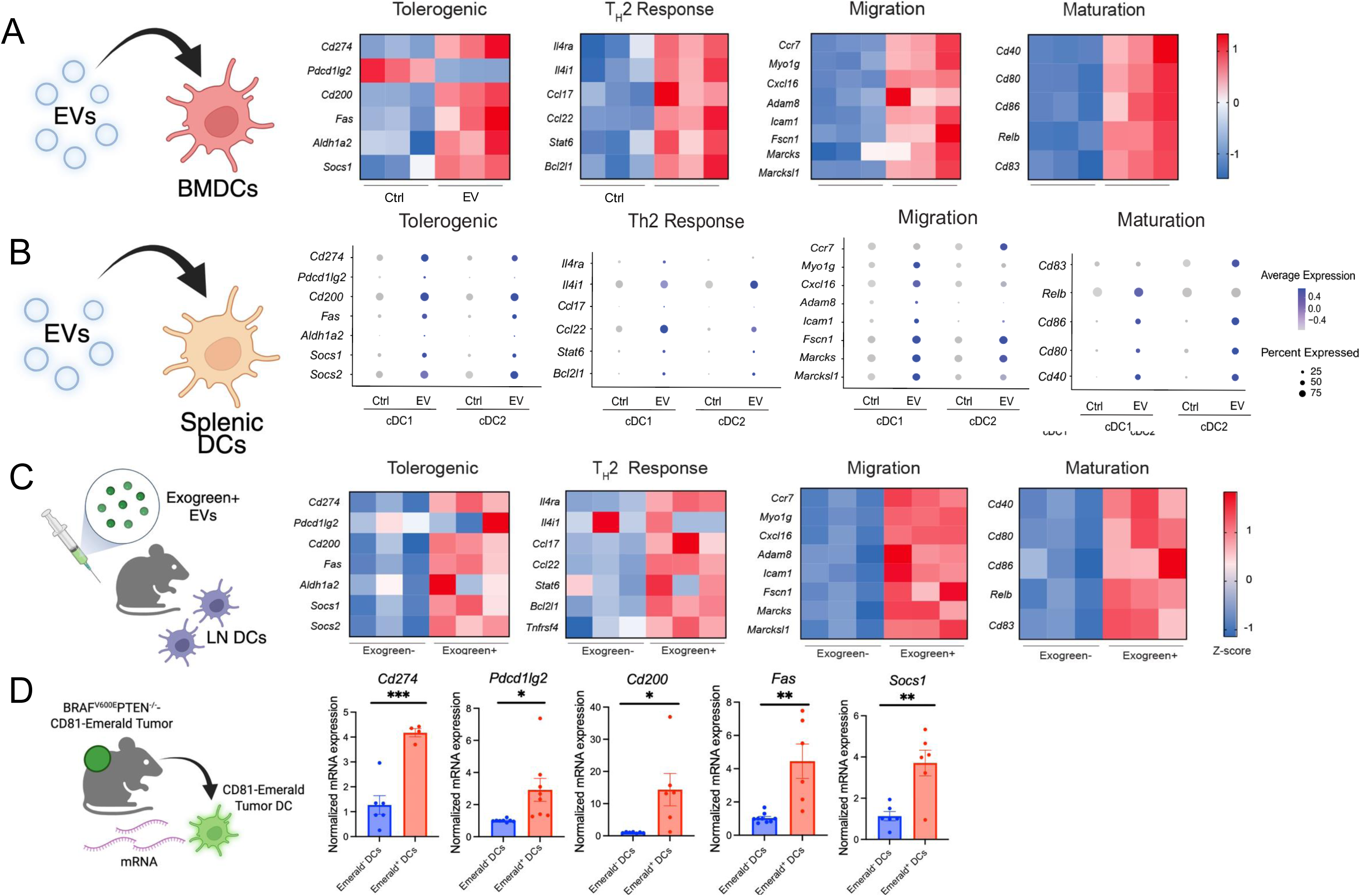
Tumor-derived EVs induce a tolerogenic transcriptional signature in DCs. **(A)** Heatmap derived from bulk RNA-sequencing (RNA-seq) analysis of bone marrow-derived DCs (BMDCs) treated with PBS (Ctrl) or tumor-derived EVs (10 µg/ml) for 24h *in vitro*. Differentially expressed genes are clustered by functional module: Tolerogenic, TH2 Response, Migration, Maturation. The color scale represents row-normalized gene expression (Z-score). Representative of two independent experiments. **(B)** Dot plot visualization of single-cell RNA sequencing (scRNA-seq) data from splenic DC subsets (cDC1 and cDC2) treated with PBS (Ctrl) or tumor-derived EVs (20 µg/ml) for 24h *in vitro*. The size of each dot reflects the percentage of cells expressing the indicated gene, while the color intensity indicates the average expression level (scaled). Representative of two independent experiments. **(C)** Transcriptional profiling of EV-uptaking DCs *in vivo*. Exogreen-labeled tumor EVs were injected into the footpads of WT mice. Draining LNs were harvested at 48–72 h, and DCs were sorted into EV-uptaking (ExoGreen^+^) and bystander (Exogreen^-^) populations by FACS. The heatmap displays the row-normalized Z-scores of differentially expressed genes between the two subsets. Representative of two independent experiments. **(D)** Relative mRNA expression of select tolerogenic genes in tumor-infiltrating DCs. DCs were sorted from BRAF^V600E^PTEN^-/-^ tumors stably expressing CD81-Emerald Green. Gene expression is compared between EV-uptaking (GFP+) and bystander (GFP-) DC subsets (n=4-8). Data is aggregated from two independent experiments. All data are reported as mean ± SEM. \**p*<0.05, \*\**p*<0.01, \*\*\**p*<0.001 by unpaired, two-tailed Student’s t-test (D). **BMDC**, bone marrow-derived dendritic cell; **cDC**, conventional dendritic cell; **EV**, extracellular vesicle; **FACS**, fluorescence-activated cell sorting; **RNA-seq**, RNA sequencing; **scRNAseq**, single-cell RNA sequencing; **GFP**, green fluorescent protein; **mRNA**, messenger RNA; **T_H_2**, T helper 2; **WT**, wild-type.

### Tumor EVs Activate an ATF4–SREBP2 Axis to Drive Lipid Metabolic Reprogramming and Induce the mregDC Phenotype

To dissect the underlying mechanism by which tumor-derived EVs reprogram DCs toward an immunoregulatory phenotype, we assessed whether EVs activate the SREBP2 metabolic pathway. Immunoblotting of nuclear extracts indeed revealed that EVs induced nuclear accumulation of SREBP2 in DCs in a pH-independent fashion, while additional studies found no clear impact of tumor EVs on DC SREBP1 nuclear translocation (**figure 3A, online supplemental figure 3A, B)**. Prior studies have identified a role for ER stress in the activation of SREBP2 and have further implicated the protein kinase RNA-like endoplasmic reticulum kinase (PERK)-activating transcription factor 4 (ATF4) ER stress pathway as a positive regulator of SREBP2 activation^58–60^. Notably, we also found that SREBP2 upregulation in DCs coincided with the activation of the upstream ER stress effector, ATF4, while no impact was observed on ATF6 (**figure 3B, online supplemental figure 3C)**. In addition, pharmacological inhibition of the upstream ER stress sensor, PERK, abrogated SREBP2 nuclear translocation (**figure 3C**), suggesting that tumor EVs induce SREBP2 activation at least partially through triggering the PERK–ATF4 pathway. To verify that tumor-derived EVs induce SREBP2 upregulation by DCs *in situ*, we performed studies using the CD81-mEmerald fusion protein-expressing BRAF^V600E^PTEN^-/-^ melanoma model and observed tumor-infiltrating mEmerald^+^ DCs also exhibit enhanced *Srebp2* expression relative to mEmerald^-^ controls, a finding consistent with SREBP2 activation (**figure 3D**).

**Figure 3.**
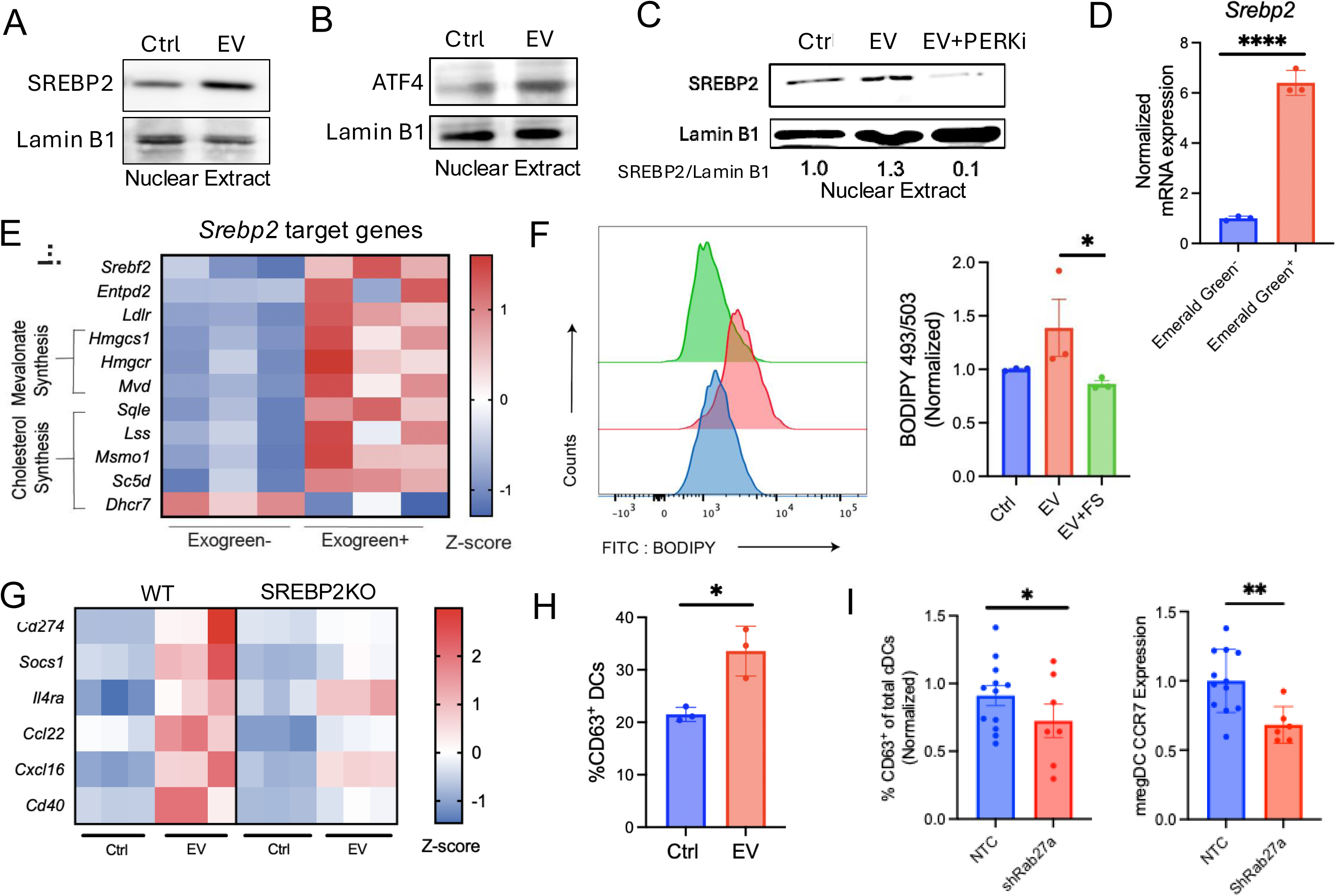
Tumor-derived EVs drive tolerogenic reprogramming and lipid accumulation via the PERK/ATF4-SREBP2 axis. (A–B) Western blot analysis of nuclear extracts from BMDCs treated with PBS (Ctrl) or tumor-derived EVs (20 µg/ml; 24h). Immunoblots show the nuclear enrichment of the mature transcription factors SREBP2 **(A)** and ATF4 **(B)**. Lamin B1 serves as the loading control for the nuclear fraction. Each representative of three independent experiments. **(C)** Western blot analysis of nuclear SREBP2 levels in BMDCs treated with PBS (Ctrl), EVs (20 µg/ml; 24h), or EVs (20 µg/ml; 24h) with a PERK inhibitor (GSK2606414; 500 nM, preincubation time 1h; 24 h). Densitometric quantification of the SREBP2 band intensity normalized to Lamin B1 band intensity performed using ImageJ. Representative of three independent experiments. **(D)** Relative *Srebp2* expression levels between EV-uptaking (Emerald Green^+^) and bystander (Emerald Green^-^) DC subsets. DCs were isolated from BRAF^V600E^PTEN^-/-^ tumors expressing CD81-Emerald Green using FACS (n=3). Representative of two independent experiments. **(F)** Flow cytometry analysis of neutral lipid accumulation in DCs. Cells were treated with PBS (Ctrl), EVs (20 µg/ml; 24h), or EVs in the presence of the SREBP2 inhibitor Fatostatin (FS). Shown are representative histograms of BODIPY 493/503 fluorescence (*left*) and the quantification of mean fluorescence intensity (MFI) normalized to the Ctrl group (*right*) (n=3). Representative of three independent experiments. **(G)** Heatmap displaying the relative mRNA expression of tolerogenic genes (*Cd274, Socs1, Il4ra, Ccl22, Cxcl16, Cd40*) in BMDCs generated from WT or DC-restricted Srebp2-knockout (KO) mice (CD11c-*Srebp2*^-/-^). Cells were treated with PBS (Ctrl) or EVs (20 µg/ml) for 24 h. The color scale represents row-normalized gene expression (Z-score). Representative of two independent experiments. **(H)** Quantification of the frequency of CD11c^+^CD63^+^ DCs in BMDC cultures treated with PBS (Ctrl) or EVs (10 µg/ml) for 24 h (n=3). Representative of two independent experiments. **(I)** Impact of tumor EVs on mregDC development *in vivo*. TDLNs were harvested from mice harboring BRAF^V600E^PTEN^-/-^-NTC or BRAF^V600E^PTEN^-/-^-shRab27a melanomas (as described in figure 1). *Left*, normalized frequency of CD11c^+^MHCII^hi^CD63^+^ cDCs. *Right*, Relative CCR7 expression by mregDCs (n=7–11). Data aggregated from two independent experiments. All data are reported as mean ± SEM. \**p*<0.05, \*\**p*<0.01, \*\*\*\**p*<0.0001. Data analyzed by unpaired, two-tailed Student’s t-test (D), Kruskal-Wallis test followed by Dunn’s multiple comparisons test (F), unpaired two-tailed Student’s t-test (H), and one-tailed Student’s t-test (I). **cDC**, conventional dendritic cell; **Ctrl**, control; **DC**, dendritic cell; **EV**, extracellular vesicle; **FACS**, fluorescence-activated cell sorting; **FS**, Fatostatin; **KO**, knockout; **MFI**, mean fluorescence intensity; **mregDC**, migratory regulatory dendritic cell; **mRNA**, messenger RNA; **NTC**, non-targeting control; **PERKi**, PERK inhibitor; **shRNA**, short hairpin RNA; **TDLN**, tumor-draining lymph node; **WT**, wild-type.

To determine whether EV-mediated SREBP2 activation also translates to functional metabolic reprogramming, we analyzed lipid metabolic pathways in EV-treated DCs. Following the intra-dermal footpad administration of Exo-Green-labeled tumor EVs, RNAseq transcriptional analysis identified a marked upregulation of known target genes of SREBP2, including several cholesterol biosynthetic genes in ExoGreen^+^ DCs isolated by fluorescence-activated cell sorting (FACS) from the draining popliteal LNs (**figure 3E**). Consistent with transcriptional activation of this metabolic pathway, BODIPY staining demonstrated that tumor EVs induced a significant increase in neutral lipid accumulation in DCs, an effect that was abrogated by pharmacologic inhibition of SREBP2 (**figure 3F**). As we have previously demonstrated SREBP2 to contribute to mregDC development, we next tested whether SREBP2 is required for tumor EV-mediated induction of the tolerogenic mregDC transcriptional program^24^. Using *Srebf2*-silenced DCs from a DC-restricted *Srebf2*^-/-^ mouse model, we found that tumor EVs were no longer able to induce the upregulation of the previously reported mregDC transcriptional signature, suggesting that SREBP2 is likely necessary for mregDC development in response to tumor-derived EVs (**figure 3G)**. We, as well as others, have demonstrated that mregDCs exhibit unique surface expression of the CD63 tetraspanin^24^ ^61^. Consistent with this finding, tumor EV exposure promotes the emergence of CD63⁺ DCs in bone marrow cultures, a phenotype further supported by RNA-seq analysis identifying *Cd63* as a top upregulated gene in LN DCs *in vivo* (**figure 3H, online supplemental figures 2D, 3D**). Conversely, the suppression of tumor EV release by silencing *Rab27a* expression correlated with diminished numbers of CD63^+^CCR7^+^ mregDCs in the TME (**figure 3I, online supplemental figure 3E**). Altogether, these results support a signaling cascade in which tumor-derived EVs induce ER stress, which in turn, drives SREBP2-dependent lipid metabolic reprogramming resulting in the differentiation of CD63⁺ mregDCs in the TME.

### Tumor EVs Induce IRE1α/XBP1s Signaling in DCs

Prior studies have observed robust ER stress and X-box binding protein 1 (XBP1) activation to accompany increased lipid stores in tumor-associated DCs in ovarian cancer ^30^ ^33^. However, the underlying mechanism governing the induction of this pathway within the TME has remained unclear. We performed additional Gene Set Enrichment Analysis (GSEA) of the previously described scRNAseq dataset derived from tumor EV-treated splenic DCs, revealing the upregulation of multiple pathways predominantly associated with oncogenic signaling, metabolic reprogramming, and cellular stress, including the unfolded protein response (UPR) pathway (**figure 4A, online supplemental figure 4A**). Inositol-requiring enzyme 1 (IRE1α) is the most conserved UPR stress sensor that functions as an endoribonuclease to process *Xbp1* mRNA, thereby generating the active spliced XBP1 isoform, XBP1s^62^. To determine whether tumor EVs may induce IRE1α-XBP1s activation in DCs, we performed Western Blot analysis probing for IRE1α phosphorylation and XBP1s production without or with the IRE1α inhibitor, 4u8c (**figures 4B,C, online supplemental figure 4B**). These data demonstrated that tumor-derived EVs induce the accumulation of spliced XBP1 (XBP1s) in DCs in an IRE1α-dependent manner (**figure 4C**). Additional studies found EVs derived from the EO771 breast cancer cell line as well as the APC^-/-^Kras^G12D^p53^-/-^SMAD4^-/-^ (AKPS) colon cancer cell line also upregulated XBP1s protein expression, while EVs derived from the NIH-3T3 fibroblast cell line did not, suggesting that EVs broadly generated by malignant cells may exhibit this unique property (**online supplemental figures 4C, D**). Consistent with this work, we also found tumor EVs to induce the expression of the XBP1s and its target genes, *Sec61a* and *Dnajb9*, in DCs based on quantitative real-time PCR analysis (**figure 4D**)^33^. To assess whether tumor EVs may trigger the activation of the IRE1α-XBP1s pathway in DCs *in vivo*, we administered Exo-Green-labeled tumor EVs by intra-dermal footpad injection as described above and interrogated FACS-sorted Exo-Green^+^ DCs by RNAseq analysis. In line with our *in vitro* findings, we observed upregulation of several XBP1s target genes (*Icam1*, *Dnajb2*, *Clip2*, *Sec61a2* and *Dnajb9*) in these isolated LN DCs (**figure 4E**)^63^ ^64^. To further verify that bona fide tumor EVs induce the IRE1α-XBP1 pathway in DCs in the context of tumor, we returned to the CD81-mEmerald-expressing BRAF^V600E^PTEN^-/-^ melanoma model and analyzed FACS-isolated mEmerald^+^ DCs for XBP1s target gene expression by qrt-PCR. This work also showed an elevation in the expression of XBP1s target genes (*Sec61a* and *Dnajb9*) relative to control mEmerald^-^ DCs isolated from BRAF^V600E^PTEN^-/-^ melanomas (**figure 4F**). Overall, these data imply that tumor EVs represent at least one trigger by which tumors can promote XBP1 activation and drive DC lipid accumulation. Notably, although prior work has implicated reactive oxygen species (ROS) as an inducer of ER stress in DCs, we found no evidence that tumor EVs were promoting lipid peroxidation, suggesting that tumor EV-induced ER stress is likely occurring through an alternative mechanism (**online supplemental figure 4E**).

**Figure 4.**
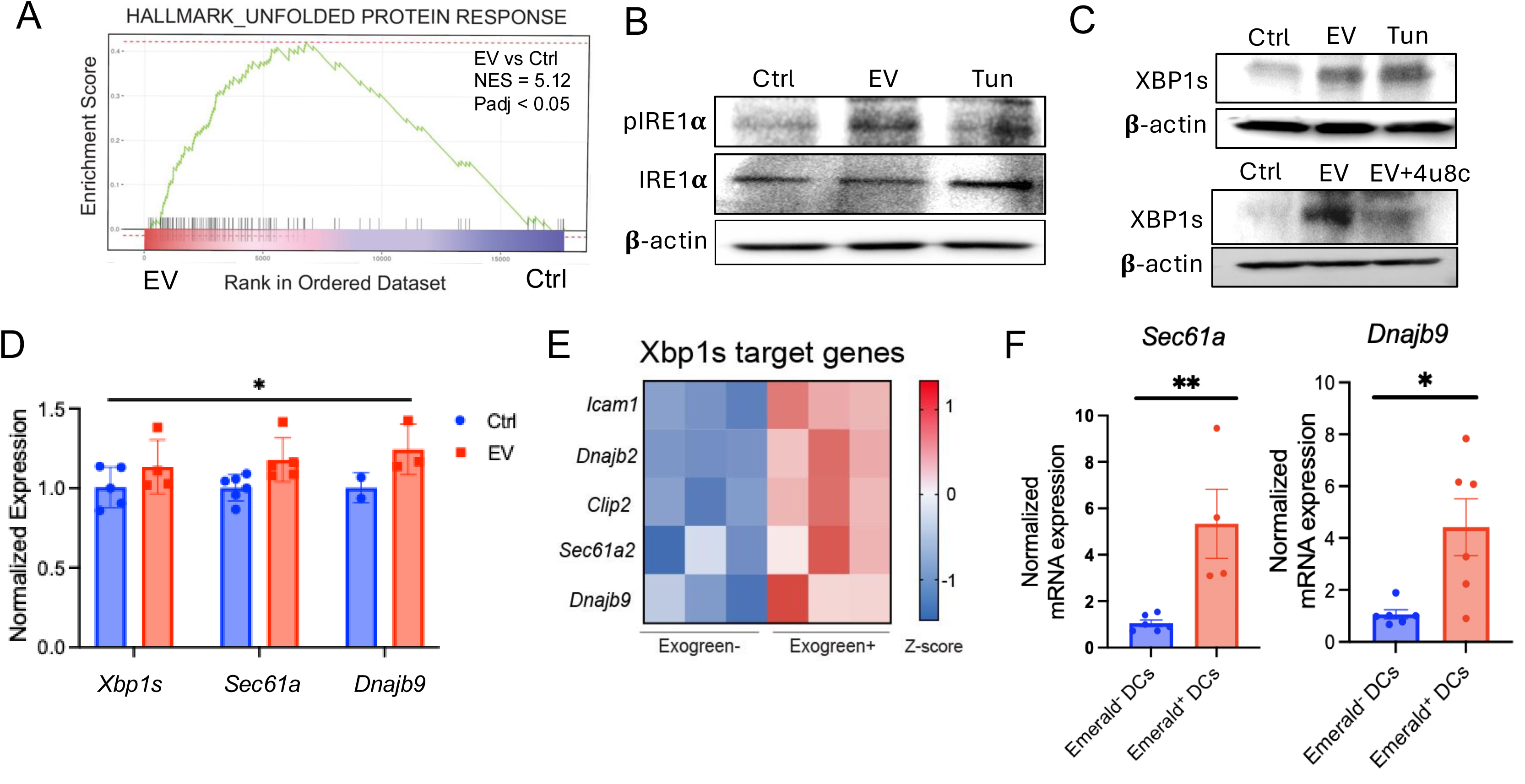
EV-induced IRE1α-XBP1s signaling in DCs. **(A**) Gene set enrichment analysis (GSEA) plot showing significant enrichment of the Hallmark Unfolded Protein Response (UPR) gene signature in EV-treated splenic DCs compared to Controls. The normalized enrichment score (NES = 5.12) and adjusted p-value (Padj < 0.05) are displayed. Representative of two independent experiments. **(B)** Western blot analysis of phosphorylated IRE1α (p-IRE1α) and total IRE1α in splenic DCs treated with EVs or tunicamycin (Tun) as a positive control. Representative of four independent experiments. **(C)** Western blot analysis of spliced XBP1 (XBP1s) protein expression in BMDCs. *Top*, Cells were treated with PBS (Ctrl), tumor-derived EVs (20 µg/ml; 24h), or Tunicamycin (Tun; 10µM; 4h) as a positive control. *Bottom*, Analysis of XBP1s levels in BMDCs treated with EVs in the presence or absence of the IRE1α inhibitor, 4µ8c. β-actin serves as the cytoplasmic loading control. Representative of four independent experiments. **(D)** Normalized mRNA expression of XBP1s and its target genes (*Sec61a, Dnajb9*) in splenic DCs treated with PBS (Ctrl) or tumor EVs (20 µg/ml; 24h) (n=3-5). **(E)** Heatmap of relative gene expression (Z-score) for XBP1s target genes (*Icam1, Dnajb2, Clip2, Sec61a2, Dnajb9*) in EV-uptaking (Exogreen^+^) versus bystander (Exogreen^-^) LNDCs isolated *in vivo*. Representative of two independent experiments. **(F)** Normalized mRNA expression of XBP1s and its target genes (*Sec61a, Dnajb9*) in tumor-infiltrating DCs, comparing EV-uptaking (Emerald Green^+^) versus bystander (Emerald Green^-^) DC subsets (n=4-5). Data is aggregated from two independent experiments. All data is reported as mean ± SEM. \**p*<0.05, \*\**p*<0.01 by two-way ANOVA (D) and unpaired Student’s t-test (F). **EVs**, extracellular vesicles; **XBP1s**, spliced X-box binding protein 1; **Tun**, tunicamycin (ER stress inducer); **BMDCs**, bone marrow-derived dendritic cells; **LNDCs**, lymph node dendritic cells; **Ctrl**, control.

### Tumor EVs Reprogram DC Lipid Metabolism through an IRE1α-XBP1s-PPAR-α Signaling Axis

Elevated lipid accumulation is a well-described characteristic of tumor-associated DC populations that has been correlated with impaired DC-dependent antigen cross-presentation and stimulation of CD8^+^ T cell responses^53^. Consistent with a role for the UPR pathway in regulating DC functionality through enhanced lipid accumulation, we found tumor EVs to increase DC neutral lipid stores in a manner dependent upon IRE1α activation (**figure 5A**). To further dissect the underlying mechanism of this metabolic shift, we returned to the transcriptional dataset derived from tumor EV-treated splenic DCs and identified an enrichment of a gene set regulated by the lipid-sensing nuclear receptor, peroxisome proliferator-activated receptor alpha (PPAR-α) (**figure 5B**). Additional experiments further indicated that tumor EVs induce PPAR-δ expression to a lesser extent than PPAR-α expression in DCs (**figure 5C, online supplemental figures 5A, B**). In addition, we found the inhibition of PPAR-α to suppress DC lipid stores induced by tumor EVs more dramatically than the inhibition of either PPAR-δ or PPAR-γ (**online supplemental figures 5C).** Previous studies have demonstrated the XBP1 pathway to induce the expression of PPAR-α ^65^. To determine if PPAR-α activation is a direct consequence of IRE1α/XBP1s signaling pathway activation in response to tumor EVs, we pharmacologically inhibited the UPR sensor IRE1α and found this was sufficient to suppress both tumor EV-mediated PPAR-α nuclear accumulation and PPAR-α DNA binding activity (**figures 5D,E**). We then sought to determine whether PPAR-α activity was required for the observed lipid accumulation in response to tumor EVs. Indeed, pharmacological inhibition of PPAR-α by the small molecule inhibitor, TPST-1120, was sufficient to inhibit the EV-induced increase in neutral lipid content, suggesting that the XBP1-PPAR-α signaling axis is a critical regulator of lipid accumulation in DCs, likely via the upregulation of lipid scavenger receptors, such as CD36 (**figure 5F, online supplemental figure 5D**). As PPAR-α is a master transcription factor that governs fatty acid oxidation (FAO), we utilized a flow cytometry-based assay to verify that tumor-derived EVs robustly stimulate FAO activity in DCs (**figure 5G**). Given our prior work establishing that enhanced FAO is a functional hallmark of tolerogenic DCs, these data collectively suggest that tumor EVs drive DC dysfunction by engaging a PPARα-dependent FAO pathway^22^. This conclusion is further corroborated *in vivo*, where tumor-infiltrating DCs that have engulfed tumor-derived EVs exhibit increased expression of the canonical PPAR-α target gene and rate-limiting FAO enzyme, *Cpt1a* (**figure 5H**). These findings suggest that tumor-derived EVs trigger an IRE1α-XBP1s-PPARα signaling axis, leading to both lipid accumulation and enhanced FAO in DCs.

**Figure 5.**
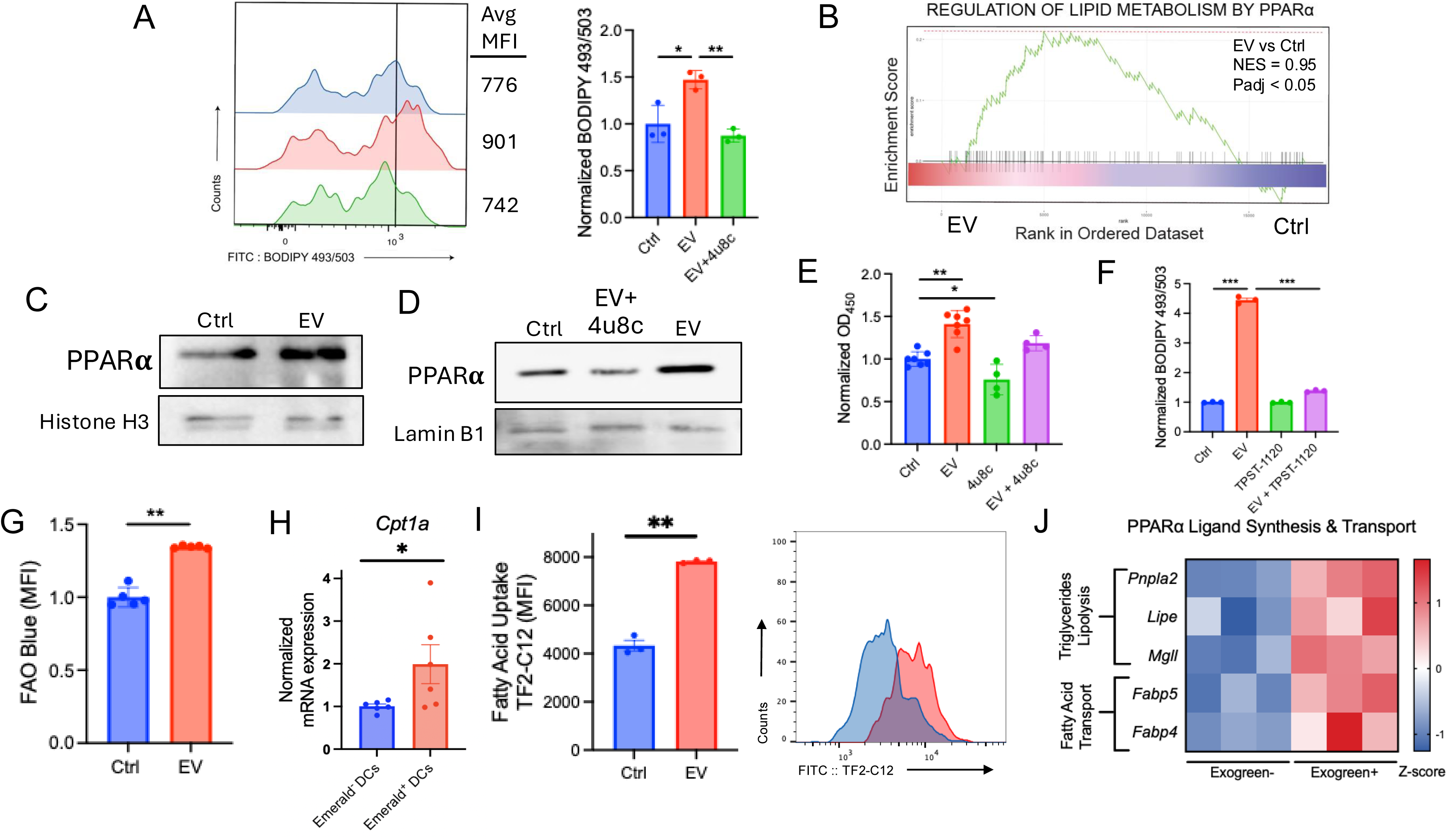
EV-induced UPR signaling drives lipid accumulation and PPAR-α activation in DCs. **(A)** Flow cytometry analysis of neutral lipid accumulation in splenic DCs treated with PBS (Ctrl), tumor EVs (20 µg/ml; 24h), or tumor EVs (20 µg/ml; 24h) in the presence of the IRE1 α inhibitor 4μ8c. *Left*, Representative histograms of BODIPY 493/503 fluorescence. *Right*, quantification of the normalized mean fluorescence intensity (MFI) (n=3). Representative of three independent experiments. **(B)** GSEA plot from the scRNAseq data in figure 2B, showing enrichment of the "Regulation of Lipid Metabolism by PPARα" gene set in EV-treated DCs compared to controls. Adjusted p-value (Padj < 0.05) is displayed. Representative of two independent experiments. **(C)** Western blot analysis of nuclear PPAR-α levels in BMDCs treated with or without EVs (20 µg/ml; 24h). Histone H3 is shown as a nuclear loading control. Representative of three independent experiments. **(D)** Immunoblot analysis of nuclear PPAR-α protein expression in BMDCs following treatment with PBS (Ctrl), tumor EVs (20 µg/ml; 24h), or EVs in the presence of the IRE1α inhibitor 4μ8c. Lamin B1 serves as the nuclear loading control. Representative of three independent experiments. **(E)** Quantification of PPAR-α DNA-binding activity in nuclear extracts from BMDCs treated with EVs (20 µg/ml; 24h), 4μ8c, or both (n=4-7). Representative of three independent experiments. **(F)** Flow cytometry quantification of neutral lipid content (BODIPY 493/503) in DCs treated with EVs (20 µg/ml; 24h) with or without the PPAR-α inhibitor TPST-1120. Representative of three independent experiments. **(G)** Flow cytometric analysis of fatty acid β-oxidation (FAO) activity in splenic DCs treated with PBS (Ctrl) or EVs (20 µg/ml; 24h) using FAO-Blue (n=5). **(H)** Normalized mRNA expression of *Cpt1a*, a key rate-limiting enzyme in FAO and a known PPAR-α target gene, in tumor-infiltrating DCs from the model in Figure 2D, comparing EV-uptaking (Emerald Green^+^) versus bystander (Emerald Green^-^) DC subsets (n=6). Representative of two independent experiments. **(I)** Heatmap of relative gene expression (Z-score) for triglyceride lipolysis (*Pnpla2*, *Lipe*, *Mgll*) and fatty acid transport genes (*Fabp5*, *Fabp4*) in EV-uptaking (Exogreen^+^) versus bystander (Exogreen^-^) LNDCs isolated *in vivo*. Representative of two independent experiments. **(J)** Flow cytometric analysis of fatty acid uptake by splenic DCs treated with PBS (Ctrl) or EVs (20 µg/ml; 24h) using TF2-C12. *Right*, representative histogram. Representative of three independent experiments. All data is reported as mean ± SEM. \**p*<0.05, \*\**p*<0.01, \*\*\**p*<0.001, \*\*\*\**p*<0.0001 based on a one-way ANOVA (A,E,F) and an unpaired Student’s t-test (G,H,J). **FAO**, fatty acid oxidation; **GSEA**, gene set enrichment analysis; **MFI**, mean fluorescence intensity; **PPARα**, peroxisome proliferator-activated receptor alpha; **BMDCs**, bone marrow-derived dendritic cells; **Ctrl**, control ; **EVs**, extracellular vesicles; **IRE1α,** inositol-requiring enzyme 1 alpha.

We were interested in understanding the observed relationship between enhanced cholesterol synthesis and increased FAO as observed in EV-exposed DCs. We previously observed that mregDCs exhibit enhanced low-density lipoprotein (LDL) receptor surface expression based on flow cytometry^24^. This is consistent with our more recent observation that EV exposure also augments LDL receptor expression in DCs (**figure 3E**). Together, these findings suggest that EV-exposed DCs take up increased levels of triglyeride-derived fatty acids, which we confirmed based on TF2-C12 flow cytometry (**figure 5I**). Indeed, transcriptional studies demonstrate an upregulation in the expression of several triglyeride lipolysis enzymes and fatty acid transporters in EV-exposed DCs, a finding that correlates with a downregulation in several fatty acid synthesis enzymes (**figure 5J, online supplemental figure 5E**). Overall, these studies suggest that SREBP2 activation drives elevated fatty acid levels; thereby, fueling FAO in tolerogenic DCs. Given our prior studies revealing a key role in FAO driving DC tolerization, these findings also suggest that FAO may represent a common pathway that is necessary for DCs to acheive a pro-tolerogenic state^22^.

### PPAR-α Signaling Contributes to the Immunosuppressive Effects of Tumor EVs

To more rigorously study the role of the PPAR-α signaling pathway in tumor EV-mediated DC tolerization, we engineered a DC-specific PPAR-α knockout mouse model by crossing mice harboring the cDC-restricted *Zbtb46* promoter-driven cre recombinase with PPAR^fl/fl^ mice^66^ (**online supplemental figures 6A,B**). We implanted both the control BRAF^V600E^PTEN^-/-^ melanoma cell line and the *Rab27*-silenced BRAF^V600E^PTEN^-/-^ melanoma cell line into these syngeneic control Zbtb46-Cre and Zbtb46-PPARα^-/-^ hosts. As expected, the genetic silencing of Rab27a in control Zbtb46-Cre mice resulted in a significant suppression of tumor growth compared to non-targeting controls (NTC) (**figure 1A**). Strikingly, this tumor-suppressive effect was diminished in DC-specific PPARα-deficient mice, suggesting that DC-intrinsic PPAR-α signaling is important for mediating the immunosuppressive properties of tumor EVs (**figure 6A**). We then utilized multi-parameter flow cytometry to determine the impact of tumor-derived EVs on the immune microenvironment. This work demonstrated that suppressed tumor EV production results in enhanced levels of tumor-infiltrating CD8^+^ T cells as well as a reduction in KLRG1^+^CD4^+^FoxP3^+^ Tregs, consistent with our prior work (**figures 1B, 6B, C, online supplemental figures 6C,D**). However, these changes were largely abrogated in hosts harboring DCs selectively silenced for *Ppara* (**figures 6B,C**). We then isolated DCs from either Zbtb46-PPARα^-/-^ or control Zbtb46-Cre mice. After confirming that *Ppara*-silencing did not compromise DC viability, we tested the impact of PPAR-α on tumor EV-dependent alterations in DC function (**online supplemental figure 6E**). While tumor EVs significantly suppressed DC-dependent CD8⁺ T cell proliferation in control hosts, this effect was partially reversed in *Ppara*-silenced DCs, (**figure 6D, online supplemental figure 6F**). In line with this finding, tumor EV stimulation of DC-mediated CD4^+^FoxP3^+^ Treg differentiation was also suppressed upon *Ppara*-silencing (**figure 6E, online supplemental figure 6G**). Overall, these results suggest that tumor EVs induce their pro-tolerogenic effects on DCs partially through PPARα signaling, a finding that is largely consistent with our prior studies emphasizing the important role of FAO in DC tolerization19 22.

**Figure 6.**
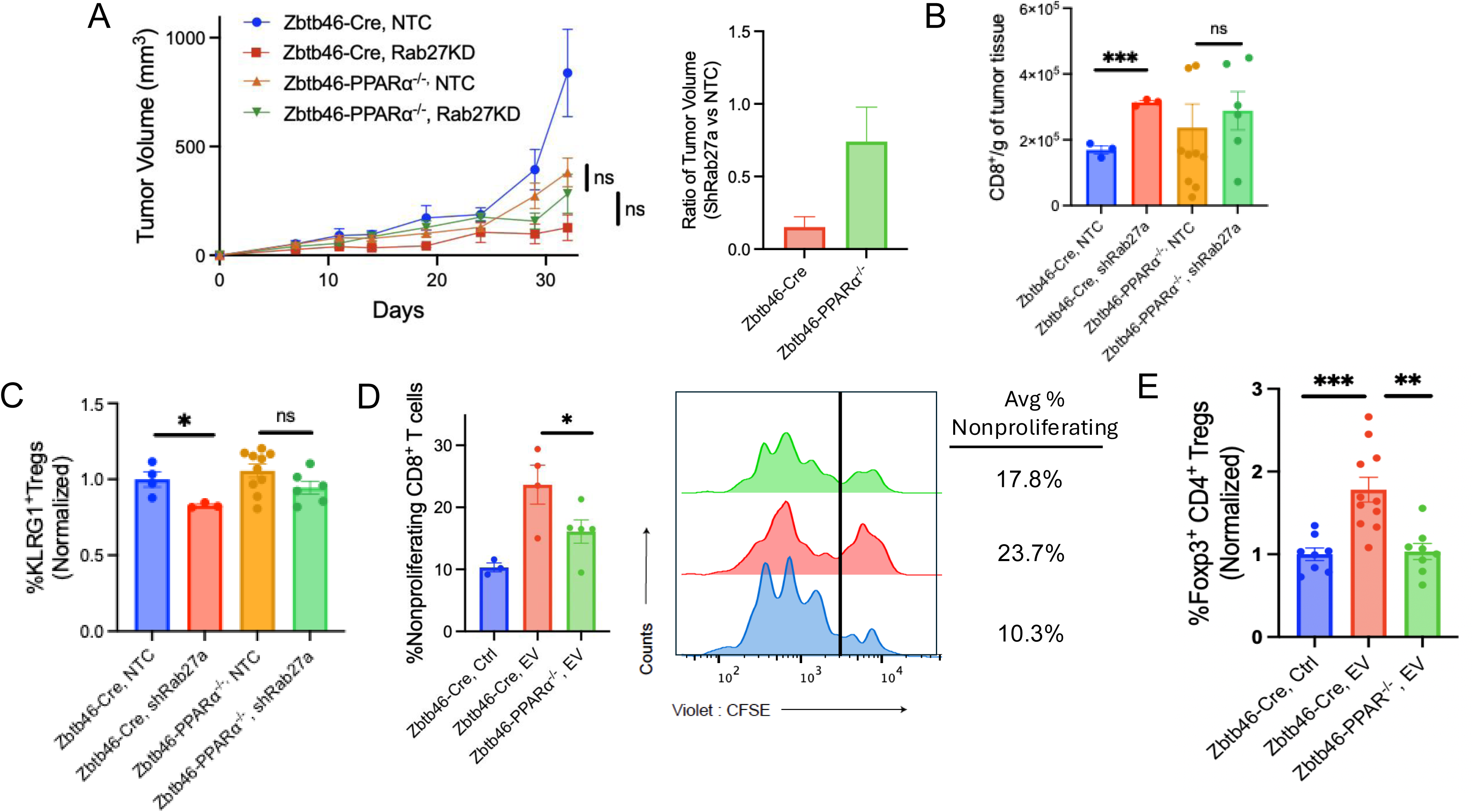
PPAR-α deficiency in DCs promotes tumor immune escape via an EV-dependent mechanism. **(A)** *In vivo* tumor growth kinetics in Zbtb46-Cre (control) and Zbtb46-*Ppara*^-/-^ mice following implantation of BRAF^V600E^PTEN^-/-^-NTC or BRAF^V600E^PTEN^-/-^-shRab27a melanomas. *Right*, The bar graph displays BRAF^V600E^PTEN^-/-^-shRab27a : BRAF^V600E^PTEN^-/-^-NTC melanoma ratios in Zbtb46-Cre (control) and Zbtb46-*Ppara*^-/-^ mice. Representative of two independent experiments. **(B)** Tumors from the experiment in **(A)** were harvested and analyzed by CD8^+^ T cell flow cytometry analysis. Absolute number of CD8^+^ T cells are normalized per gram of tumor (n=5-12 mice/group). Representative of two independent experiments. **(C)** Tumors from the experiment in **(A)** were harvested and analyzed by KLRG1^+^CD4^+^FoxP3^+^ T_reg_ flow cytometry analysis (n=5-12 mice/group). Representative of two independent experiments. **(D)** *In vitro* CD8^+^ T cell proliferation assay. Splenic DCs isolated from the Zbtb46-Cre (control) and Zbtb46-*Ppara*^-/-^ mice were treated with PBS (Ctrl) or EVs (20 µg/ml; 24h) and then co-cultured with CFSE-labeled naive CD8^+^ T cells for 60 hours. Non-proliferating (CFSE^high^) CD8^+^ T cells are quantified. *Right*, Representative histograms are shown (n=3-5). Representative of two independent experiments. **(E)** *In vitro* Treg differentiation assay. Splenic DCs from Zbtb46-Cre or Zbtb46-*Ppara*^-/-^ mice were treated with PBS (Ctrl) or tumor EVs (20 µg/ml; 24h) and co-cultured with naive CD4^+^ T cells. FoxP3^+^CD4^+^ Tregs were quantified by flow cytometry (n=8-12). Data is aggregated from two independent experiments. All data is reported as mean ± SEM. \**p*<0.05, \*\**p*<0.01, \*\*\**p*<0.001 based on one-way ANOVA (B-E). **CFSE**, carboxyfluorescein succinimidyl ester; **DC**, dendritic cell; **KLRG1**, killer cell lectin-like receptor G1; **NTC**, non-targeting control; **PPARα**, peroxisome proliferator-activated receptor alpha; **Ctrl**, control; **EVs**, extracellular vesicles; **FoxP3**, forkhead box P3; **ns**, non-significant; **shRNA**, short hairpin RNA.

### Pharmacologic Inhibition of PPAR-α Synergizes with Anti-PD-1 to Suppress Tumor Growth and Metastasis

The oral PPAR-α inhibitor, TPST-1120, has been shown to augment the efficacy of the standard-of-care regimen, atezolizumab/bevacizumab, for the first line treatment of patients with unresectable or metastatic hepatocellular carcinoma in a phase Ib/II clinical trial^67^. More recent studies have further shown TPST-1120 to overcome anti-PD-1 resistance in select patient populations with advanced solid tumors in a phase I clinical trial^68^. Based on these data as well as our pre-clinical findings indicating that tumor-derived EVs drive an immunosuppressive DC phenotype via a PPARα-dependent pathway, we hypothesized that pharmacological inhibition of PPAR-α could overcome immune tolerance and potentiate the efficacy of immune checkpoint blockade. To test this, we initially performed a study in a syngeneic model of BRAF^V600E^PTEN^-/-^ melanoma that showed the addition of the PPAR-α antagonist, TPST-1120, to anti-PD-1 immunotherapy to nearly eliminate tumor progression (**online supplemental figure 7A**). Given these results, we then treated tumor-bearing autochthonous BRAF^V600E^PTEN^-/-^ mice with TPST-1120 alone or in combination with an anti-PD-1 antibody. While neither TPST-1120 monotherapy nor anti-PD-1 antibody monotherapy had no impact on tumor progression, the TPST-1120/anti-PD-1 antibody combination therapy resulted in synergistic suppression of tumor growth (**figure 7A**). Consistent with its effects on primary tumor growth, TPST-1120/anti-PD-1 antibody combination therapy also showed a trend towards a reduction in pulmonary metastasis as quantified by expression of the melanoma-associated TRP2 antigen in lung tissues (**figure 7B**). We further characterized the immunologic response to TPST-1120/anti-PD-1 antibody combination therapy by multi-parameter flow cytometry and found combination therapy to augment CD8⁺ T cell infiltration while concurrently suppressing intratumoral levels of CD4^+^FoxP3^+^ Tregs (**figures 7C, D, online supplemental figures 7B, C**). We then investigated whether these observed immunological changes were associated with the metabolic reprogramming of DCs in the TME. To assess the metabolic state of tumor-infiltrating conventional type 1 DCs (cDC1s), we performed SCENITH analysis, noting PPAR-α inhibition to moderately reduce the mitochondrial capacity of DC1s within the TME (**online supplemental figure 7D**)^48^. Interestingly, we found that anti-PD-1 monotherapy drove a significant increase in FAO capacity of both cDC1s and mregDCs, an effect that was eliminated by the addition of the PPAR-α antagonist TPST-1120 (**figures 7E,F**). These findings also correlated with a trend toward a reduction in the neutral lipid content of DCs within the TME following TPST-1120 treatment (**online supplemental figure 7E**). In summary, pharmacological inhibition of PPAR-α synergizes with anti-PD-1 checkpoint blockade by reversing DC FAO and enhancing anti-tumor immunity.

**Figure 7.**
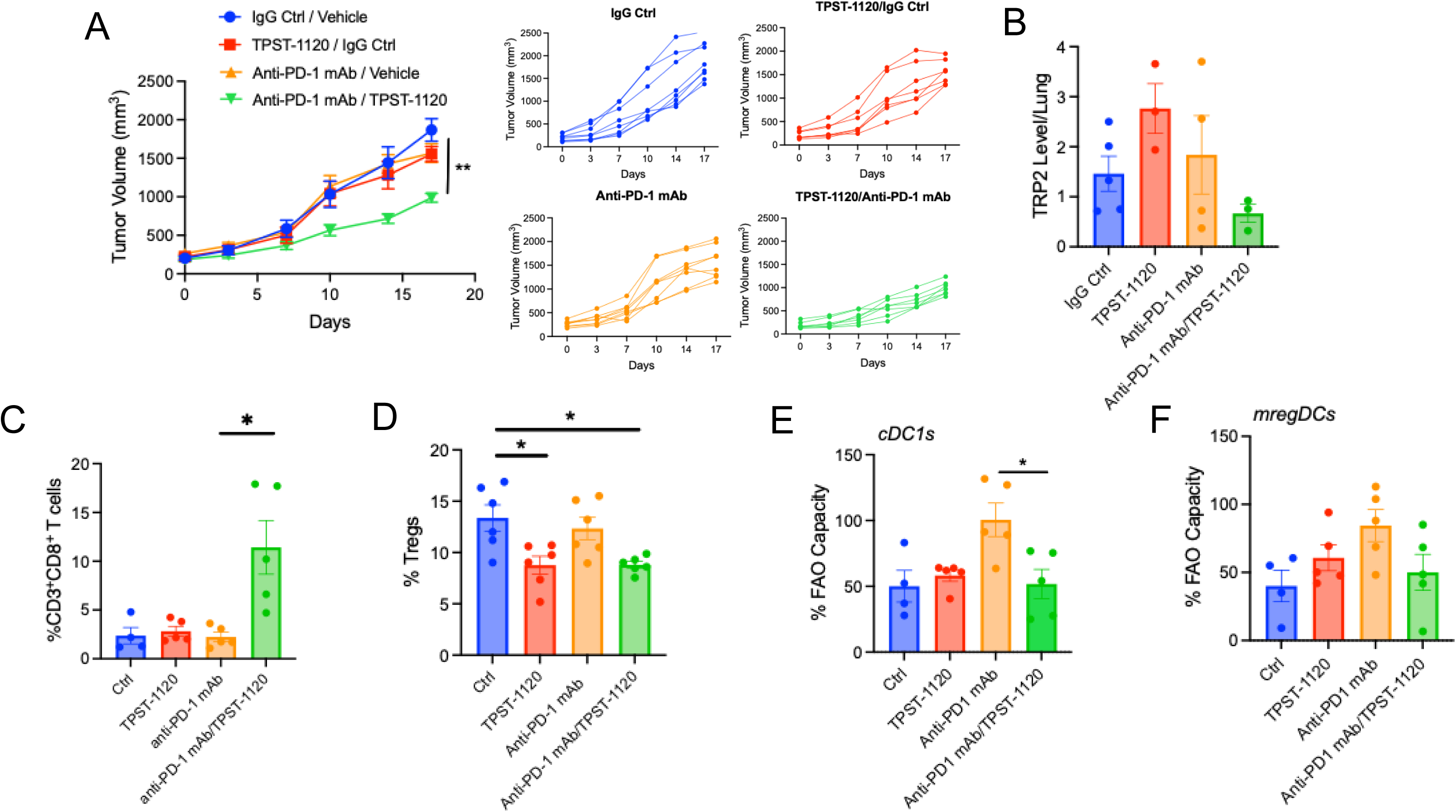
PPAR-α inhibition enhances anti-PD-1 immunotherapy in an autochthonous melanoma model. **(A)** *In vivo* efficacy of PPARα inhibition alone or in combination with anti-PD-1 immunotherapy in an autochthonous BRAF^V600E^PTEN^-/-^ melanoma model. *Right*, Spider plots showing individual tumor trajectories for each treatment group (n=5-8 mice/group). Representative of two independent experiments. **(B)** Quantification of lung metastatic burden based on qrt-PCR analysis of TRP2 levels (n=3-5). Representative of two independent experiments. **(C)** Flow cytometry analysis of tumor-infiltrating CD8^+^ T cells from the experiment described in (A), shown as a percentage of total CD3^+^ T cells (n=4-5). Representative of two independent experiments. **(D)** Flow cytometry analysis of tumor-infiltrating CD4^+^FoxP3^+^ Tregs, shown as a percentage of total CD4^+^ T cells (n=6). Representative of two independent experiments. **(E-F)** Metabolic profiling of tumor-infiltrating DCs using the SCENITH assay. The fatty acid oxidation (FAO) capacity of FACS-sorted **(E)** cDC1s and **(F)** mregDCs was assessed in tumor-bearing mice from the experiment described in (A) (n=4-5). Representative of two independent experiments. All data is reported as mean ± SEM. \**p*<0.05, \*\**p*<0.01 based on a two-way ANOVA. **FAO**, fatty acid oxidation; **mregDC**, migratory regulatory dendritic cell; **cDC1s**, conventional type 1 dendritic cells; **4-HT**, 4-hydroxytamoxifen; **mAb**, monoclonal antibody; **PD-1**, programmed cell death protein 1; **PPARα**, peroxisome proliferator-activated receptor alpha; **SCENITH**, single cell energetic metabolism by profiling translation inhibition; **TPST-1120**, PPAR-α inhibitor; **Tregs**, regulatory T cells; **TRP2**, tyrosinase-related protein.

## DISCUSSION

Given the critical role of the DC in generating anti-tumor immunity, it is likely that tumor evolution involves the development of a plethora of mechanisms that suppress their function. We are interested in exploring these pathways and determining whether common elements exist between them for future therapeutic targeting. One shared characteristic that has been well documented for dysfunctional DCs in the tumor microenvironment (TME) is increased lipid stores, alluding to an underlying metabolic derangement that is contributing to their functional impairment. In the present study, we expand on the role that tumor-derived EVs have in contributing to DC tolerization and evasion of the host immune system. This work details how tumor-derived EVs induce an ER stress response, resulting in the concurrent activation of the PERK-ATF4 and IRE1α-XBP1s UPR pathways in DCs. This coordinated signaling orchestrates metabolic reprogramming characterized by simultaneous cholesterol biosynthesis and the induction of fatty acid oxidation (FAO). This unique metabolic profile promotes intracellular lipid accumulation while enforcing a FAO-dependent metabolic program that drives an immunosuppressive DC phenotype^53^. Collectively, these findings support a hierarchical model of ER stress and metabolic regulation elicited by EVs that ultimately lead to immune evasion.

Our findings refined the current conceptual framework of how metabolic stress shapes immune function. We previously established that tumor-derived lactate promotes the development of mregDCs via the activation of the transcription factor, SREBP2. Our current data validates SREBP2 as a conserved metabolic cornerstone for DC tolerogenicity and also revealed a critical mechanistic divergence induced by tumor EVs. Unlike lactate, which functions predominantly as a local metabolic signal, tumor EVs can act as a long-range systemic stress signal. This distinct property initiates a coordinated metabolic program simultaneously enforcing SREBP2-mediated anabolism and engaging the IRE1α-XBP1s axis to transcriptionally upregulate PPARα-driven catabolism. By inducing such a stress response that couples lipid accumulation with FAO in DCs, our study demonstrates that tumors may utilize multiple immunosuppressive mechanisms that enforce an integrated state of immune tolerance.

To place our findings in context, we first address the foundational work of Cubillos-Ruiz et al. (2015), which established the IRE1α-XBP1s axis as a driver of DC dysfunction^33^. Their study demonstrated that constitutive XBP1s activation forces DCs into a state of aberrant lipid biosynthesis, characterized by the upregulation of lipogenic enzymes and the accumulation of immunosuppressive triglycerides. Our data confirmed this fundamental link between ER stress and lipid dysregulation and expanded upon these findings by demonstrating that tumor EVs serve as an upstream environmental trigger of this pathway. More recently, Yin et al. (2020) demonstrated that tumor EVs suppress DC function via the activation of the PPAR-α transcription factor^69^. Our work confirms these findings while also identifying the proximal signaling mechanism required for inducing PPAR-α transcriptional activity. However, while Yin et al. (2020) focused on passive lipid availability, our data suggests a distinct, active mechanism of intracellular lipid mobilization. Specifically, we observed the robust upregulation of genes governing triglyceride lipolysis (*Pnpla2, Lipe, Mgll*) and fatty acid transport (*Fabp4, Fabp5*), concurrent with the downregulation of *de novo* fatty acid synthesis machinery. Together, these preliminary data suggests that EVs drive the mobilization of intracellular lipid reservoirs. The convergence of SREBP2-mediated cholesterol synthesis and enhanced fatty acid uptake provides an abundance of substrates that maximizes the basal esterification capacity, thereby driving lipid droplet formation in DC populations within the TME. This metabolic rewiring maintains a reservoir of fatty acid substrates required to sustain PPARα-driven FAO, while simultaneously preserving a pool of free cholesterol that may support the mature phenotype of tolerogenic DCs. Mechanistically, the simultaneous engagement of cholesterol mobilization and FAO explains the functional paradox of the mregDC transcriptional state. Belabed et al. (2025) established that NPC1-mediated cholesterol mobilization is a prerequisite for assembling the lipid nanodomains that stabilize CCR7 and MHC-II on the DC surface^70^. Consistent with this prerequisite, the upregulation of *Npc2* in EV-recipient LNDCs indicates active cholesterol mobilization may support the conserved migratory capacity and mature surface phenotype observed in mregDCs. However, our study uncovers a critical metabolic divergence that dictates the functional outcome of this maturation. While Belabed et al. (2025) demonstrate that cholesterol mobilization is essential for establishing the competency for inflammatory signaling, we demonstrate that this potential is compromised in the TME. Indeed, the concurrent induction of FAO establishes a dominant tolerogenic metabolic program that overrides the pro-inflammatory effects typically driven by cholesterol mobilization. Thus, while cholesterol mobilization provides the structural basis for maturation and migration, it is the metabolic reprogramming via FAO that ultimately drives these cells into a tolerogenic state, preventing them from initiating an effective anti-tumor immune response. These findings offer further support that FAO serves as a metabolic rheostat for modulating immunity. Whereas pharmacological inhibition of this pathway may contribute to overcoming immune tolerance in the TME, its stimulation could be conversely leveraged to restore peripheral tolerance in autoimmune disorders.

We must acknowledge the limitations of this work. While syngeneic tumor models lack the immune sculpting observed during the evolution of autochthonous tumor systems, they represent the current limit of technical feasibility for dissecting many aspects of tumor EV function *in vivo*. Crucially, this model does enable the monitoring of endogenous fluorescently-labeled EV transfer within a fully competent immune microenvironment. However, the development of additional autochthonous experimental models to examine the impact of tumor EVs on tumor immunity will be an important step toward understanding the role of EVs in tumor immune evasion. We attempted to ensure that the quantity of EVs utilized in our studies reflected physiologic levels in the TME, however it is difficult to know exactly what EV levels individual tumors are exposed to at any given location or at any point in time. Due to this uncertainty, it will be important to develop methods to study the impact of tumor-derived EVs on anti-tumor immune responses in patients. This study also did not address the specific properties of tumor-derived EVs responsible for inducing UPR activation in DCs. Whether this effect is secondary to the unique lipid components of tumor-derived EVs or a factor harbored by tumor-derived EVs remains unclear and is in need of further study.

Tumor EVs have largely been ignored in terms of immunotherapy development, however the current study serves to highlight their importance in tumor-mediated immune evasion. Given that several stimuli seem to rely on the induction of similar metabolic pathways to promote DC tolerization in the TME, the targeting of common downstream signaling nodes may serve as promising strategies to overcome the immunotherapy resistance that is observed more commonly in the clinic. Determining which of these stimuli is more active for any particular tumor to enable the rationale tailoring of therapeutics will be another challenge that will need to be addressed in future studies.

## ACKNOWLEDGEMENTS

The authors would like to thank Tempest Therapeutics, Inc. for providing the PPAR-α inhibitor, TPST-1120, for these studies. We are grateful to K. Howell for technical assistance with generating the DC-specific *Ppar*α-deficient mice and A. Hanks for assisting with the BRAF^V600E^PTEN^-/-^-shRab27a melanoma study. We would also like to thank Dr. Donald P. McDonnell (Duke University) for generously providing the EO771 cell line and Dr. Jatin Roper (Duke University) for providing the AKPS cell line. Finally, we would also like to express our thanks to Dr. Frank A. Gomez (National Cancer Institute) for kindly providing the Pparα^fl/fl^ mice.

## CONTRIBUTORS

BH and MP developed the original hypotheses and designed the experiments. XW, MS, and MP performed the experiments and analyzed the data. YN and MB provided technical suppor for all studies. BH, XW, and MP wrote and edited the manuscript. XW, MP, MB, MS, BT, and BH assisted with manuscript review. BH supervised the study.

