## Supplementary Material for "Tumor-derived Extracellular Vesicles Induce ER Stress to Drive Tolerogenic Dendritic Cell Development in the Tumor Microenvironment"

Figure S1.

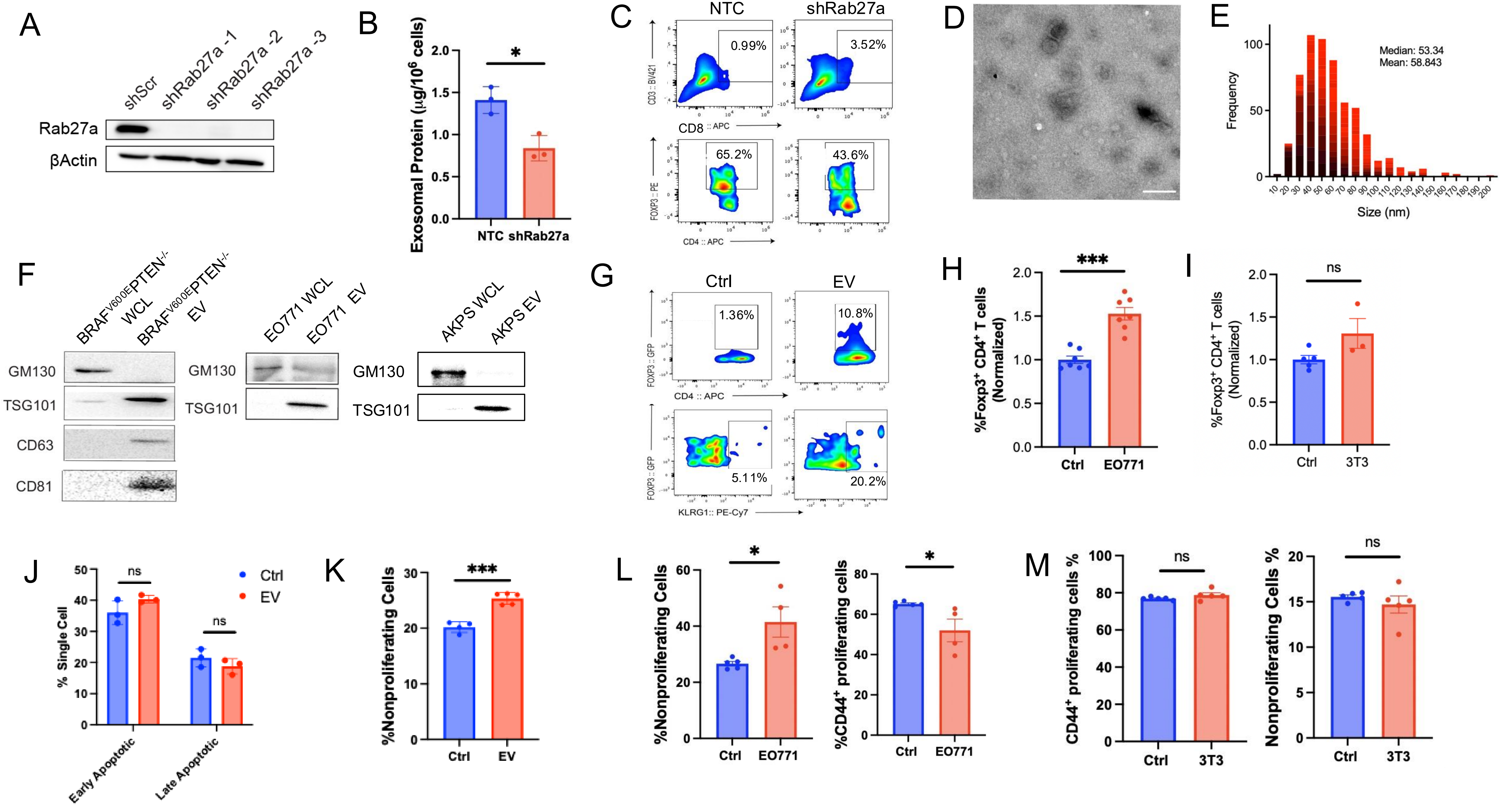
A
shScr
shRab27a -1
shRab27a -2
shRab27a -3
Rab27a
βActin
B
Exosomal Protein (µg/10^6 cells)
*
NTC
shRab27a
C
NTC
shRab27a
0.99%
3.52%
CD3 :: BV421
CD8 :: APC
65.2%
43.6%
FOXP3 :: PE
CD4 :: APC
D
E
Median: 53.34
Mean: 58.843
Frequency
Size (nm)
F
BRAFV600EPTEN-/- WCL
BRAFV600EPTEN-/- EV
EO771 WCL
EO771 EV
AKPS WCL
AKPS EV
GM130
TSG101
CD63
CD81
G
Ctrl
EV
1.36%
10.8%
FOXP3 :: GFP
CD4 :: APC
5.11%
20.2%
KLRG1:: PE-Cy7
H
***
%Foxp3+ CD4+ T cells (Normalized)
Ctrl
EO771
I
ns
%Foxp3+ CD4+ T cells (Normalized)
Ctrl
3T3
J
ns
ns
Ctrl
EV
% Single Cell
Early Apoptotic
Late Apoptotic
K
***
%Nonproliferating Cells
Ctrl
EV
L
*
%Nonproliferating Cells
Ctrl
EO771
*
%CD44+ proliferating cells
Ctrl
EO771
M
ns
CD44+ proliferating cells %
Ctrl
3T3
ns
Nonproliferating Cells %
Ctrl
3T3

**Figure S1. Characterization of Tumor-derived EVs and Validation of Rab27a Knockdown. (A)** Western blot analysis of Rab27a expression in $BRAF^{V600E}PTEN^{-/-}$ melanoma cells transduced with non-targeting control (shScr) or three distinct shRNAs targeting Rab27a (shRab27a). β-Actin served as the loading control. Representative of three independent experiments. **(B)** Quantification of EV protein yield from the supernatants of $BRAF^{V600E}PTEN^{-/-}$-NTC and $BRAF^{V600E}PTEN^{-/-}$-shRab27a cell lines. Representative of two independent experiments. **(C)** Representative flow cytometry plots of tumor-infiltrating $CD8^+$ and $CD4^+FoxP3^+$ T cell populations from the $BRAF^{V600E}PTEN^{-/-}$-NTC and $BRAF^{V600E}PTEN^{-/-}$-shRab27a tumors in figure 1. Representative of two independent experiments. **(D)** Representative transmission electron microscopy (TEM) image of purified tumor-derived EVs. Scale bar, 100 nm. Representative of two independent experiments. **(E)** Nanoparticle tracking analysis (NTA) histogram depicting the size distribution and frequency of isolated EVs. **(F)** Western blot characterization of EVs isolated from $BRAF^{V600E}PTEN^{-/-}$ (melanoma), EO771 (breast cancer), and AKPS (colorectal cancer) cell lines. Whole cell lysates (WCL) and purified EVs were probed for the exosome markers TSG101, CD63, and CD81, and the negative control cis-Golgi marker, GM130. Representative of two independent experiments. **(G)** Representative flow cytometry plots of $FoxP3^+CD4^+$ T cells generated following the co-incubation of allogeneic naïve $CD4^+$ T cells with DCs treated with PBS (Ctrl) or tumor-derived EVs (*in vitro* Treg assays). Representative of two independent experiments. **(H-I)** Quantification of DC-mediated $FoxP3^+CD4^+$ T cell differentiation by DCs treated by **(H)** EO771 tumor-derived EVs or **(I)** NIH-3T3 fibroblast-derived EVs. **(J)** Viability analysis of EV-treated DCs. Frequencies shown of early apoptotic ($AnnexinV^+7\text{-}AAD^-$) and late apoptotic ($AnnexinV^+7\text{-}AAD^+$) cells. **(K)** Flow cytometry-based quantification of $CD8^+$ T cell proliferation following OT-1 CD8+ T cell co-culture with DCs pulsed with OVA prior to treatment with tumor-derived EVs (to examine impact of tumor EVs on OVA uptake). Percentage of non-proliferating $CFSE^{high}$ $CD8^+$ T cells shown. **(L)** Quantification of $CD8^+$ T cell proliferation following co-culture with DCs treated with PBS (Ctrl) or EO771-derived EVs. Percentage of non-proliferating ($CFSE^{high}$) and proliferating ($CD44^+CFSE^{low}$) $CD8^+$ T cells shown. **(M)** Quantification of $CD8^+$ T cell proliferation in response to DCs treated with NIH-3T3 fibroblast-derived EVs. All data are reported as mean ± SEM. *$p$<0.05, ***$p$<0.001 by unpaired Student's t-test (B, H-M). **AKPS,** $APC^{-/-}KRAS^{G12D}p53^{-/-}SMAD4^{-/-}$; **Ctrl,** control; **DC,** dendritic cell; **EV,** extracellular vesicle; **ns**, non-significant; **NTC**, non-targeting control; **TEM,** transmission electron microscopy; **WCL,** whole cell lysate**.**

Figure S2.

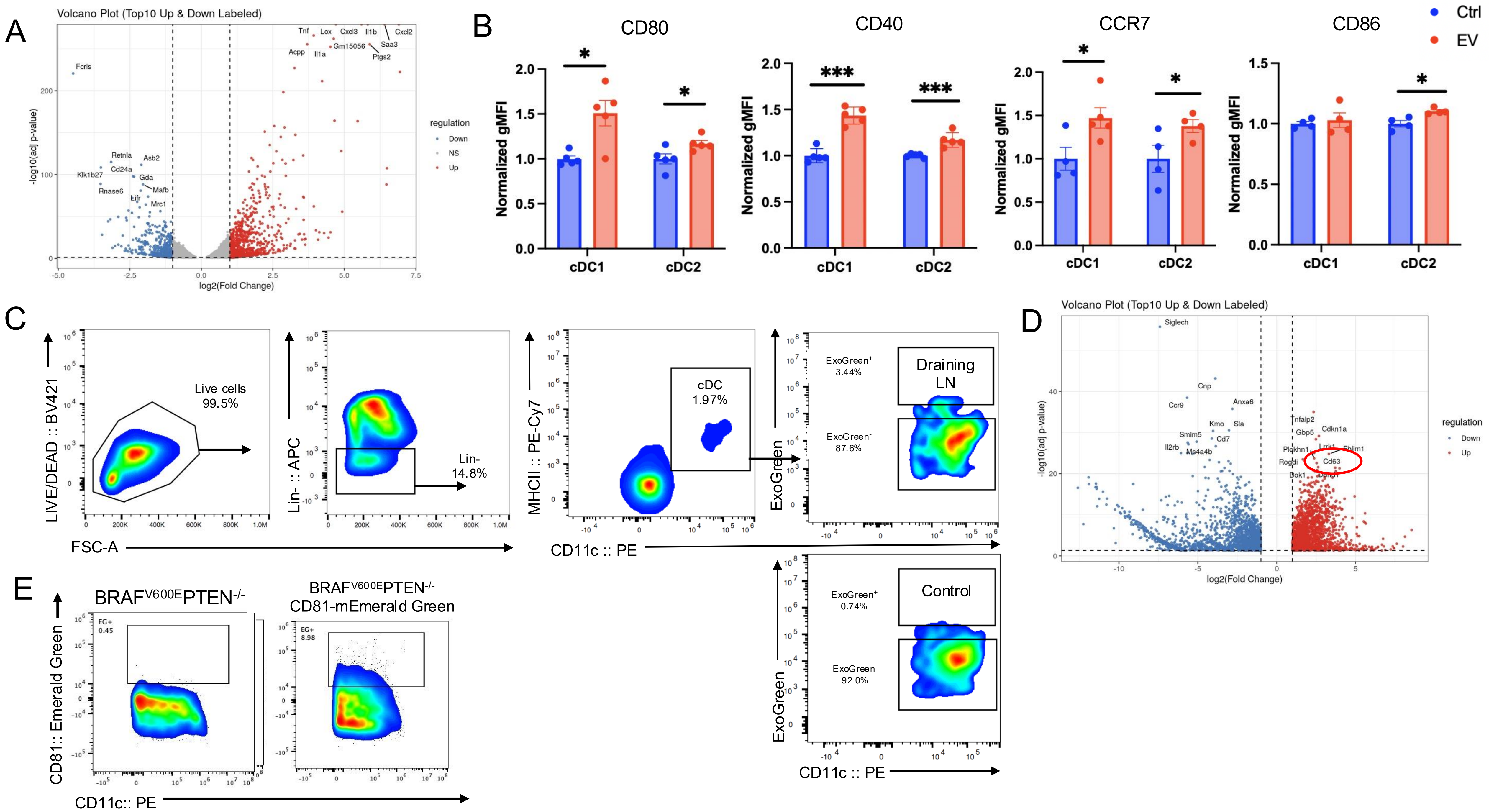
A
Volcano Plot (Top10 Up & Down Labeled)
-log10(adj p-value)
log2(Fold Change)
regulation
Down
NS
Up
B
CD80
CD40
CCR7
CD86
Ctrl
EV
Normalized gMFI
cDC1
cDC2
C
Live cells 99.5%
Lin- 14.8%
cDC 1.97%
Draining LN
ExoGreen+ 3.44%
ExoGreen- 87.6%
Control
ExoGreen+ 0.74%
ExoGreen- 92.0%
LIVE/DEAD :: BV421
Lin- :: APC
MHCII :: PE-Cy7
ExoGreen
FSC-A
CD11c :: PE
D
Volcano Plot (Top10 Up & Down Labeled)
E
BRAFV600EPTEN-/-
BRAFV600EPTEN-/- CD81-mEmerald Green
EG+ 0.45
EG+ 8.98
CD81:: Emerald Green
CD11c:: PE

**Figure S2. Transcriptional and Flow Cytometry Profiling of Dendritic Cells Following Exposure to Tumor-derived EVs. (A**) Volcano plot derived from bulk RNA-sequencing of BMDCs treated with tumor-derived EVs versus PBS (Ctrl), as detailed in figure 2A. Red dots indicate significantly upregulated genes (e.g. *Il1b, Ptgs2, Cxcl3*), while blue dots indicate significantly downregulated genes (e.g. *Fcrls*, *Retnla*, *Cd24a*). **(B)** Flow cytometry quantification of maturation markers (CD80, CD40, CD86, CCR7) on cDC1 and cDC2 subsets. Data are shown as normalized geometric mean fluorescence intensity (gMFI). Representative of two independent experiments. **(C)** Gating strategy used to identify EV-uptaking LNDCs *in vivo*. Plots show the identification of Live, $Lineage^{-}MHCII^{+}CD11c^{+}$ DCs in the TDLN and the detection of the EV marker, ExoGreen, compared to a non-draining LN control. Representative of four independent experiments. **(D)** Volcano plot displaying differential gene expression from ultra-low input RNA-sequencing of EV-uptaking ($ExoGreen^{+}$) LNDCs, corresponding to the analysis in figure 2C. The red circle highlights the upregulation of the mregDC marker, *Cd63,* in the EV-recipient population. Representative of four independent experiments. **(E)** Flow cytometry plots showing $CD81\text{-}mEmerald^{+}$ DCs following co-incubation with either control $BRAF^{V600E}PTEN^{-/-}$ melanoma cells or the CD81-mEmerald-expressing $BRAF^{V600E}PTEN^{-/-}$ melanoma cell line. CD81, EV marker. Representative of eight independent experiments. All data are reported as mean ± SEM. *$p<0.05$, ***$p<0.001$ by unpaired Student's t-test (B). **BMDC**, bone marrow-derived dendritic cell; **cDC**, conventional dendritic cell; **Ctrl**, control; **EV**, extracellular vesicle; **gMFI**, geometric mean fluorescence intensity; **Lin**, lineage; **LN**, lymph node; **LNDC**, lymph node dendritic cell**; MHCII**, major histocompatibility complex class II.

Figure S3.

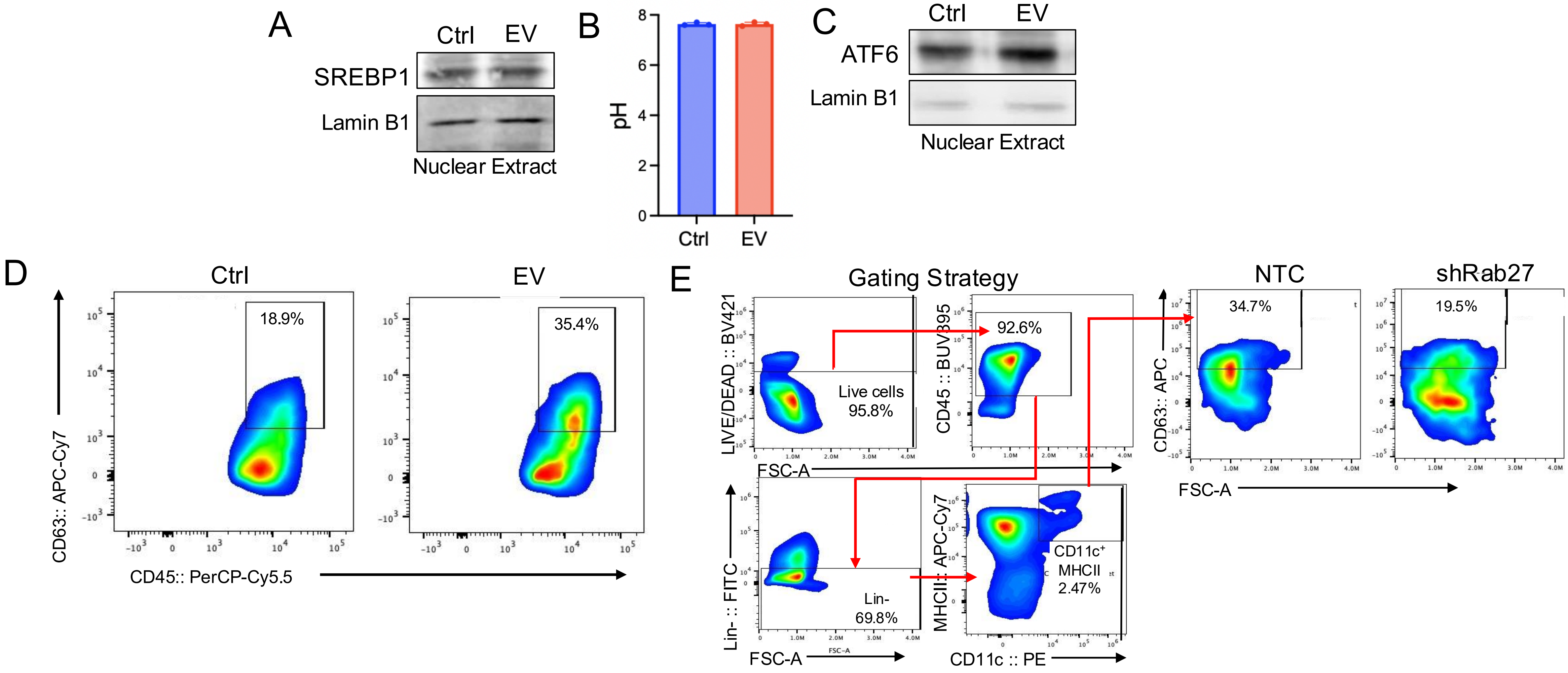
A
Ctrl
EV
SREBP1
Lamin B1
Nuclear Extract
B
pH
Ctrl
EV
C
Ctrl
EV
ATF6
Lamin B1
Nuclear Extract
D
Ctrl
EV
18.9%
35.4%
CD63:: APC-Cy7
CD45:: PerCP-Cy5.5
E
Gating Strategy
LIVE/DEAD :: BV421
Live cells
95.8%
CD45 :: BUV395
92.6%
FSC-A
Lin- :: FITC
Lin-
69.8%
MHCII :: APC-Cy7
CD11c+
MHCII
2.47%
CD11c :: PE
NTC
shRab27
34.7%
19.5%
CD63:: APC

**Figure S3. Evaluation of Alternative DC Signaling Pathways Triggered by Tumor-derived EVs. (A**) Western blot analysis of SREBP1 expression in nuclear extracts from BMDCs treated with Ctrl or tumor-derived EVs. Lamin B1 was used as a nuclear loading control. Representative of two independent experiments. **(B)** Quantification of pH levels in the culture supernatant following BMDC treatment with either PBS (Ctrl) or tumor-derived EVs. Data indicates that tumor-derived EV treatment is unlikely to trigger SREBP2 activation by lowering pH. Representative of two independent experiments. **(C)** Western blot analysis of nuclear extracts probing for the UPR sensor ATF6 to assess activation of alternative UPR branches. Lamin B1 served as the loading control. Representative of two independent experiments. **Representative Flow Cytometry Plots of DCs Following Tumor EV Exposure or Following DC Isolation from $BRAF^{V600E}PTEN^{-/-}$-NTC and $BRAF^{V600E}PTEN^{-/-}$-shRab27a Tumors (D)** Flow cytometry plots showing CD63 expression by $CD45^+$ immune cells following incubation with tumor-derived EVs compared to control. Representative of two independent experiments. **(E)** Representative flow cytometry gating strategy describing the quantification of $CD63^+$ mregDCs shown in figure 3I. DCs were identified as $Live^+CD45^+Lineage^-MHCII^+CD11c^+$ cells. Representative plots (right) display CD63 surface expression levels by cDCs isolated from $BRAF^{V600E}PTEN^{-/-}$-NTC and $BRAF^{V600E}PTEN^{-/-}$-shRab27a tumors. Representative of two independent experiments. All data are reported as mean ± SEM. **BMDC**, bone marrow-derived dendritic cell; **cDC**, conventional dendritic cell**; Ctrl**, control; **EV**, extracellular vesicle; **NTC**, non-targeting control; **shRNA**, short hairpin RNA.

Figure S4.

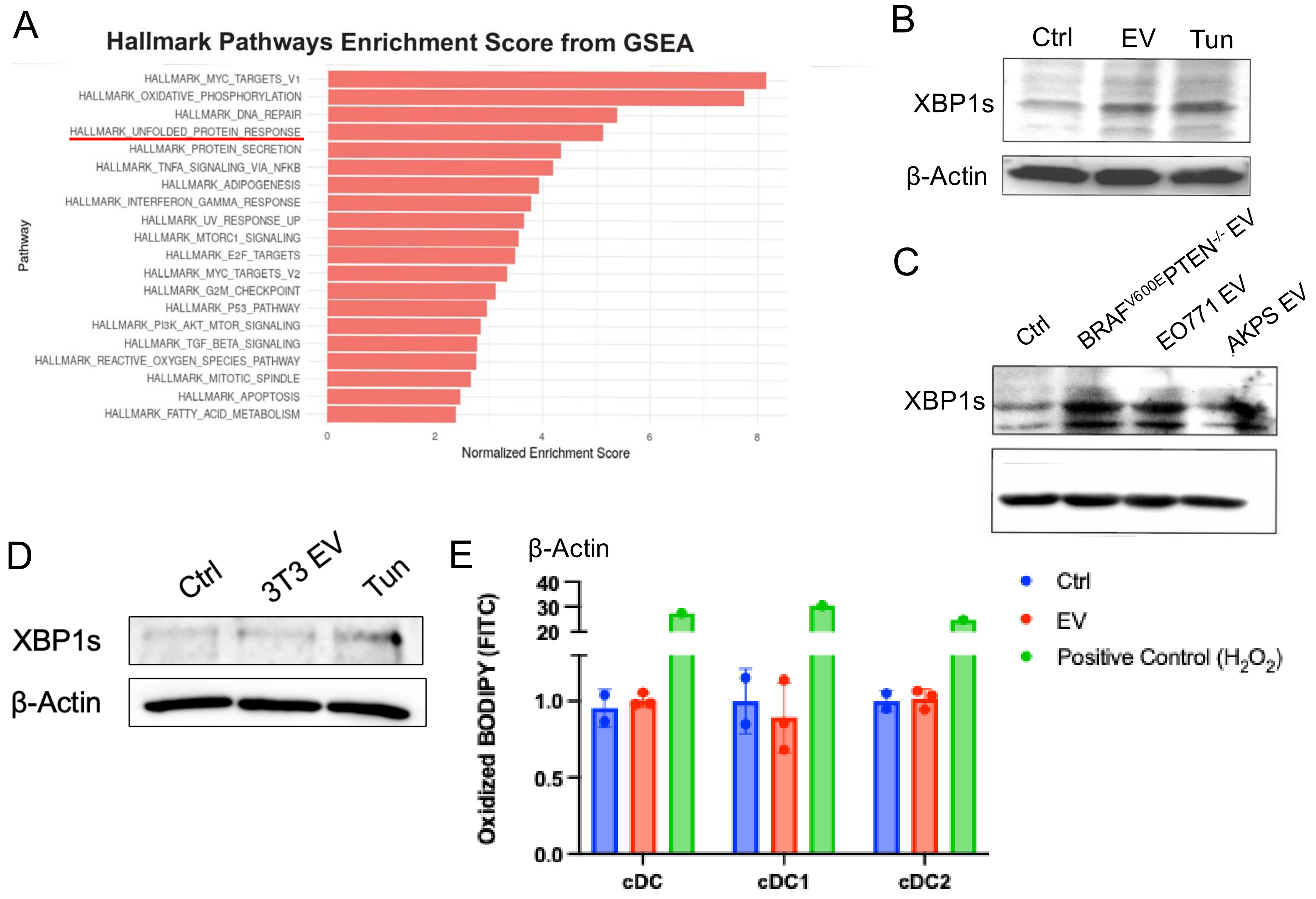
A
Hallmark Pathways Enrichment Score from GSEA
HALLMARK_MYC_TARGETS_V1
HALLMARK_OXIDATIVE_PHOSPHORYLATION
HALLMARK_DNA_REPAIR
HALLMARK_UNFOLDED_PROTEIN_RESPONSE
HALLMARK_PROTEIN_SECRETION
HALLMARK_TNFA_SIGNALING_VIA_NFKB
HALLMARK_ADIPOGENESIS
HALLMARK_INTERFERON_GAMMA_RESPONSE
HALLMARK_UV_RESPONSE_UP
HALLMARK_MTORC1_SIGNALING
HALLMARK_E2F_TARGETS
HALLMARK_MYC_TARGETS_V2
HALLMARK_G2M_CHECKPOINT
HALLMARK_P53_PATHWAY
HALLMARK_PI3K_AKT_MTOR_SIGNALING
HALLMARK_TGF_BETA_SIGNALING
HALLMARK_REACTIVE_OXYGEN_SPECIES_PATHWAY
HALLMARK_MITOTIC_SPINDLE
HALLMARK_APOPTOSIS
HALLMARK_FATTY_ACID_METABOLISM
Pathway
0
2
4
6
8
Normalized Enrichment Score
B
Ctrl
EV
Tun
XBP1s
β-Actin
C
Ctrl
BRAF$^{V600E}$PTEN$^{-/-}$ EV
EO771 EV
AKPS EV
XBP1s
β-Actin
D
Ctrl
3T3 EV
Tun
XBP1s
β-Actin
E
Oxidized BODIPY (FITC)
40
30
20
1.0
0.5
0.0
cDC
cDC1
cDC2
Ctrl
EV
Positive Control ($H_2O_2$)

**Figure S4. Tumor-derived EVs Trigger the UPR in Dendritic Cells without Altering Oxidized Lipid Levels.** **(A**) Gene Set Enrichment Analysis (GSEA) of the splenic DC cluster from the single-cell RNA sequencing dataset (corresponding to figure 2B), identifying "Hallmark Unfolded Protein Response" (highlighted) as a top enriched pathway. **(B)** Western blot analysis of XBP1s expression in BMDCs treated with PBS (Ctrl), melanoma EVs, or the ER stress inducer Tunicamycin (Tun, positive control). β-Actin served as the loading control. Representative of three independent experiments. **(C)** Western blot analysis of XBP1s in BMDCs treated with EVs isolated from $BRAF^{V600E}PTEN^{-/-}$ melanoma, EO771 breast cancer, and AKPS colorectal cancer cell lines. Representative of two independent experiments. **(D)** Western blot analysis of XBP1s in splenic DCs treated with vehicle, EVs derived from non-tumorigenic NIH/3T3 fibroblasts (3T3 EV), or Tunicamycin (Tun). Representative of two independent experiments. **(E)** Flow cytometry quantification of lipid peroxidation using BODIPYC11 probe (Oxizdized BODIPYC11/FITC) in cDC subsets. Hydrogen peroxide was used as a positive control ($H_2O_2$). Representative of two independent experiments. All data are presented as mean ± SEM. **BMDC**, bone marrow-derived dendritic cell; **cDC**, conventional dendritic cell. **Ctrl**, vehicle control; **EV**, extracellular vesicle; **GSEA**, gene set enrichment analysis; **Tun**, tunicamycin; **XBP1s**, spliced X-box binding protein 1.

Figure S5.

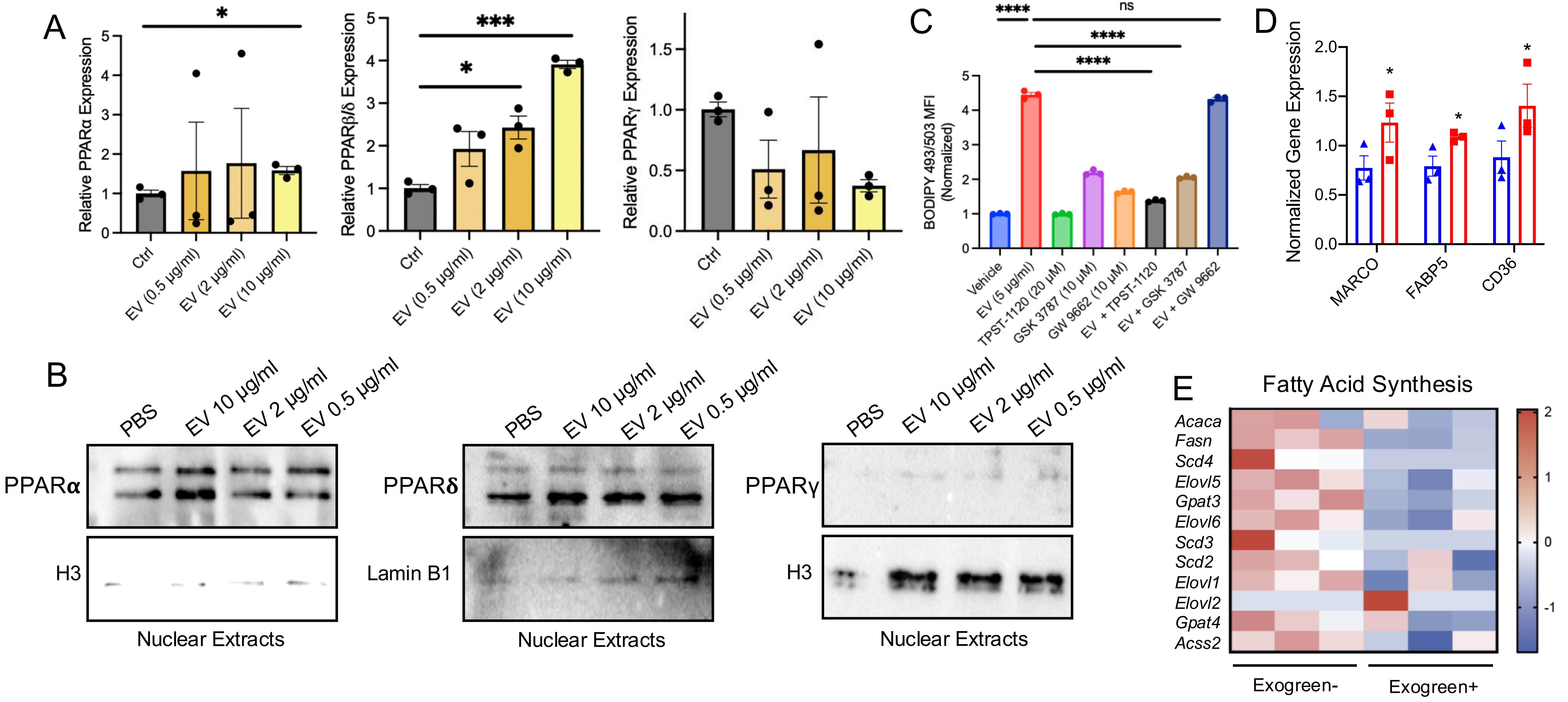
A
Relative PPARα Expression
Ctrl
EV (0.5 µg/ml)
EV (2 µg/ml)
EV (10 µg/ml)
Relative PPARβ/δ Expression
Relative PPARγ Expression
C
BODIPY 493/503 MFI (Normalized)
Vehicle
EV (5 µg/ml)
TPST-1120 (20 µM)
GSK 3787 (10 µM)
GW 9662 (10 µM)
EV + TPST-1120
EV + GSK 3787
EV + GW 9662
ns
D
Normalized Gene Expression
MARCO
FABP5
CD36
B
PBS
EV 10 µg/ml
EV 2 µg/ml
EV 0.5 µg/ml
PPARα
H3
PPARδ
Lamin B1
PPARγ
Nuclear Extracts
E
Fatty Acid Synthesis
Acaca
Fasn
Scd4
Elovl5
Gpat3
Elovl6
Scd3
Scd2
Elovl1
Elovl2
Gpat4
Acss2
Exogreen-
Exogreen+

**Figure S5. Tumor-derived EVs Upregulate PPARα Expression while Suppressing *de novo* Fatty Acid Synthesis. (A)** Quantitative real-time PCR (qrt-PCR) analysis of *Ppara, Ppard, and Ppa*rg mRNA expression in BMDCs treated with increasing concentrations of tumor-derived EVs (0.5, 2, and 10 µg/ml). Data represent relative expression normalized to control. Representative of two independent experiments. **(B)** Western blot analysis of PPAR-α, PPAR-β, and PPAR-γ expression in BMDCs treated with increasing doses of EVs. Histone H3 (H3) and Lamin B1 were used as nuclear loading controls. Representative of two independent experiments. **(C)** Flow cytometry quantification of neutral lipid accumulation (BODIPY 493/503 MFI) in DCs treated with melanoma EVs in the presence or absence of selective PPAR antagonists: TPST-1120 (PPAR-α inhibitor), GSK 3787 (PPAR-β inhibitor), and GW 9662 (PPAR-γ inhibitor). Representative of two independent experiments. **(D)** *Marco*, *Fabp5*, and *Cd36* mRNA expression levels based on qrt-PCR analysis in tumor-infiltrating DCs, comparing EV-uptaking (Emerald $Green^+$) versus bystander (Emerald $Green^-$) DC subsets. **(E)** Heatmap displaying the differential expression of genes involved in fatty acid synthesis (e.g., *Fasn, Acaca*) in EV-uptaking ($ExoGreen^+$) versus non-uptaking ($ExoGreen^-$) LNDCs, derived from the ultra-low input RNA-seq dataset shown in figure 2C. All data are reported as mean ± SEM. *$p$<0.05, ***$p$<0.001, ****$p$<0.0001 by one-way ANOVA (A, C) and unpaired Student's t-test (D). **BMDC**, bone marrow-derived dendritic cell; **Ctrl**, control; **EV**, extracellular vesicle; **LNDC**, lymph node dendritic cell; **MFI**, mean fluorescence intensity; **PPAR**, peroxisome proliferator-activated receptor.

Figure S6.

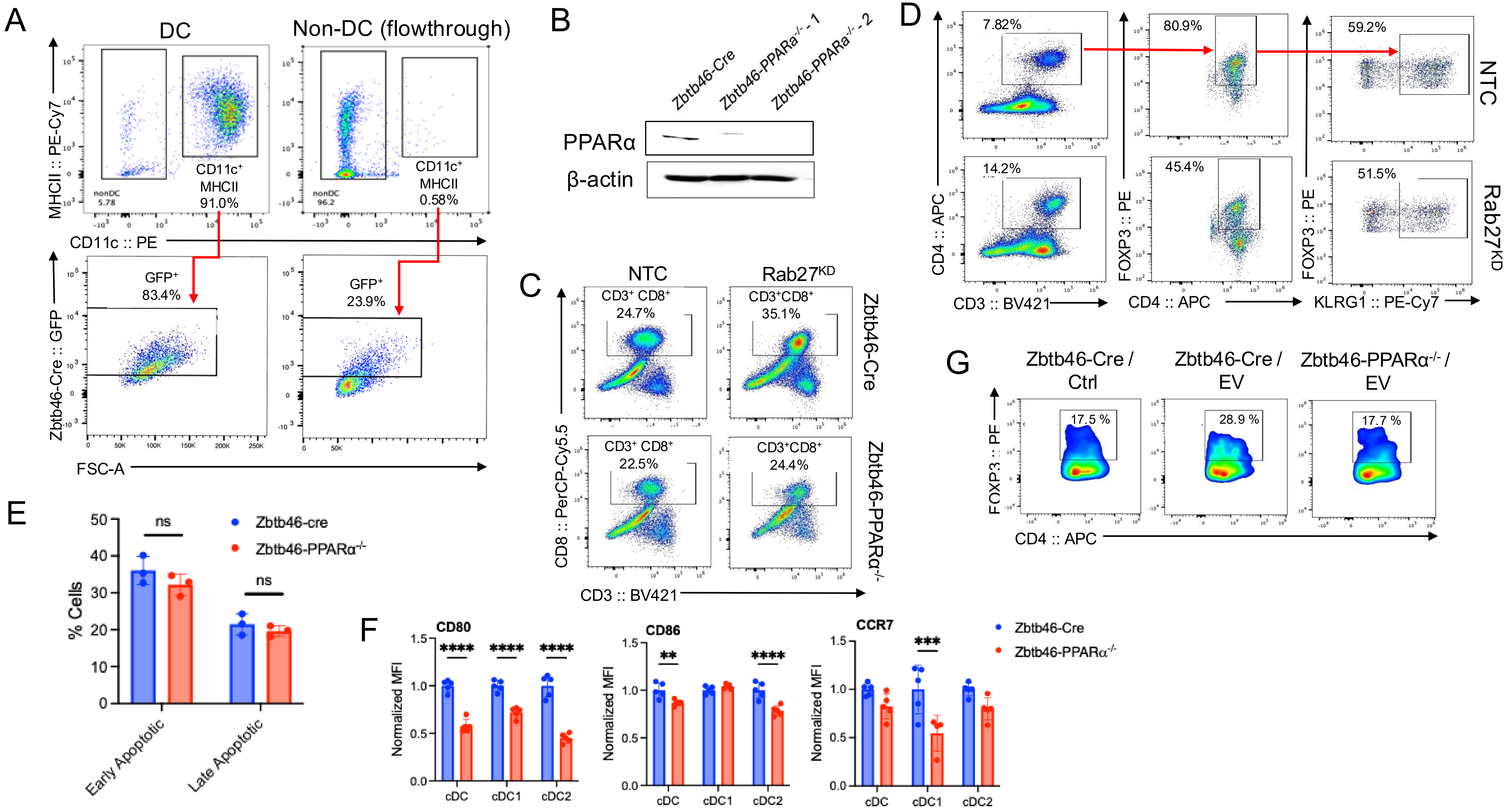

A
DC
Non-DC (flowthrough)
MHCII :: PE-Cy7
nonDC 5.78
CD11c+ MHCII 91.0%
nonDC 96.2
CD11c+ MHCII 0.58%
CD11c :: PE
GFP+ 83.4%
GFP+ 23.9%
Zbtb46-Cre :: GFP
FSC-A
B
Zbtb46-Cre
Zbtb46-PPARα-/- -1
Zbtb46-PPARα-/- -2
PPARα
β-actin
C
NTC
Rab27KD
CD3+ CD8+ 24.7%
CD3+CD8+ 35.1%
CD3+ CD8+ 22.5%
CD3+CD8+ 24.4%
Zbtb46-Cre
Zbtb46-PPARα-/-
CD8 :: PerCP-Cy5.5
CD3 :: BV421
D
7.82%
80.9%
59.2%
14.2%
45.4%
51.5%
NTC
Rab27KD
CD4 :: APC
FOXP3 :: PE
CD3 :: BV421
CD4 :: APC
KLRG1 :: PE-Cy7
G
Zbtb46-Cre / Ctrl
Zbtb46-Cre / EV
Zbtb46-PPARα-/- / EV
17.5 %
28.9 %
17.7 %
FOXP3 :: PE
CD4 :: APC
E
ns
ns
Zbtb46-cre
Zbtb46-PPARα-/-
% Cells
Early Apoptotic
Late Apoptotic
F
CD80
CD86
CCR7
Normalized MFI
cDC
cDC1
cDC2
Zbtb46-Cre
Zbtb46-PPARα-/-

**Figure S6. Validation of Dendritic Cell-specific *Ppara* Deletion and Characterization of Immune Phenotypes. (A)** Flow cytometry analysis of GFP expression in the DC ($CD11c^{+}MHCII^{+}$) and non-DC fractions of splenocytes from *Zbtb46*-Cre-GFP reporter mice. Representative of two independent experiments. **(B)** Western blot analysis of PPAR-α protein levels in splenic DCs generated from Zbtb46-Cre control and Zbtb46-PPARα$^{-/-}$ mice. Note that splenic DC purity ranges from 85-93% following CD11c microbead isolation. β-Actin was used as a loading control. Representative of three independent experiments. **(C)** Representative flow cytometry plots of tumor-infiltrating $CD8^{+}$ T cells ($CD3^{+}CD8^{+}$) in Zbtb46-Cre control or Zbtb46-PPARα$^{-/-}$ mice bearing BRAF$^{V600E}$PTEN$^{-/-}$-NTC and BRAF$^{V600E}$PTEN$^{-/-}$-shRab27a tumors. Representative of two independent experiments. **(D)** Gating strategy for tumor-infiltrating KLRG1$^{+}$CD4$^{+}$FoxP3$^{+}$ regulatory T cells in BRAF$^{V600E}$PTEN$^{-/-}$-NTC and BRAF$^{V600E}$PTEN$^{-/-}$-shRab27a tumors. Representative of two independent experiments. **(E)** Quantification of cell viability in *Zbtb46*-Cre and Zbtb46-PPARα$^{-/-}$ splenic DCs assessed by Annexin V and 7-AAD staining. Frequencies of early apoptotic (AnnexinV$^{+}$7-AAD$^{-}$) and late apoptotic (AnnexinV$^{+}$7-AAD$^{+}$) cells shown. Representative of two independent experiments. **(F)** Flow cytometry quantification of surface maturation marker expression (CD80, CD86, CCR7) on cDC, cDC1, and cDC2 subsets from Zbtb46-Cre and Zbtb46-PPARα$^{-/-}$ mice. Representative of two independent experiments. **(G)** Representative flow cytometry plots of intracellular FoxP3 expression in CD4$^{+}$ T cells following co-culture with Zbtb46-Cre or Zbtb46-PPARα$^{-/-}$ splenic DCs treated with vehicle (Ctrl) or melanoma EVs. Representative of two independent experiments. All data are reported as mean ± SEM. *$p$<0.05, **$p$<0.01, ***$p$<0.001, ****$p$<0.0001 by two-way ANOVA (E-F). **BMDC**, bone marrow-derived dendritic cell; **cDC**, conventional dendritic cell; **Ctrl**, control; **DC**, dendritic cell; **EV**, extracellular vesicle; **GFP**, green fluorescent protein; ; **KD**, knockdown; **MFI**, mean fluorescence intensity; **NTC**, non-targeting control.

Figure S7.

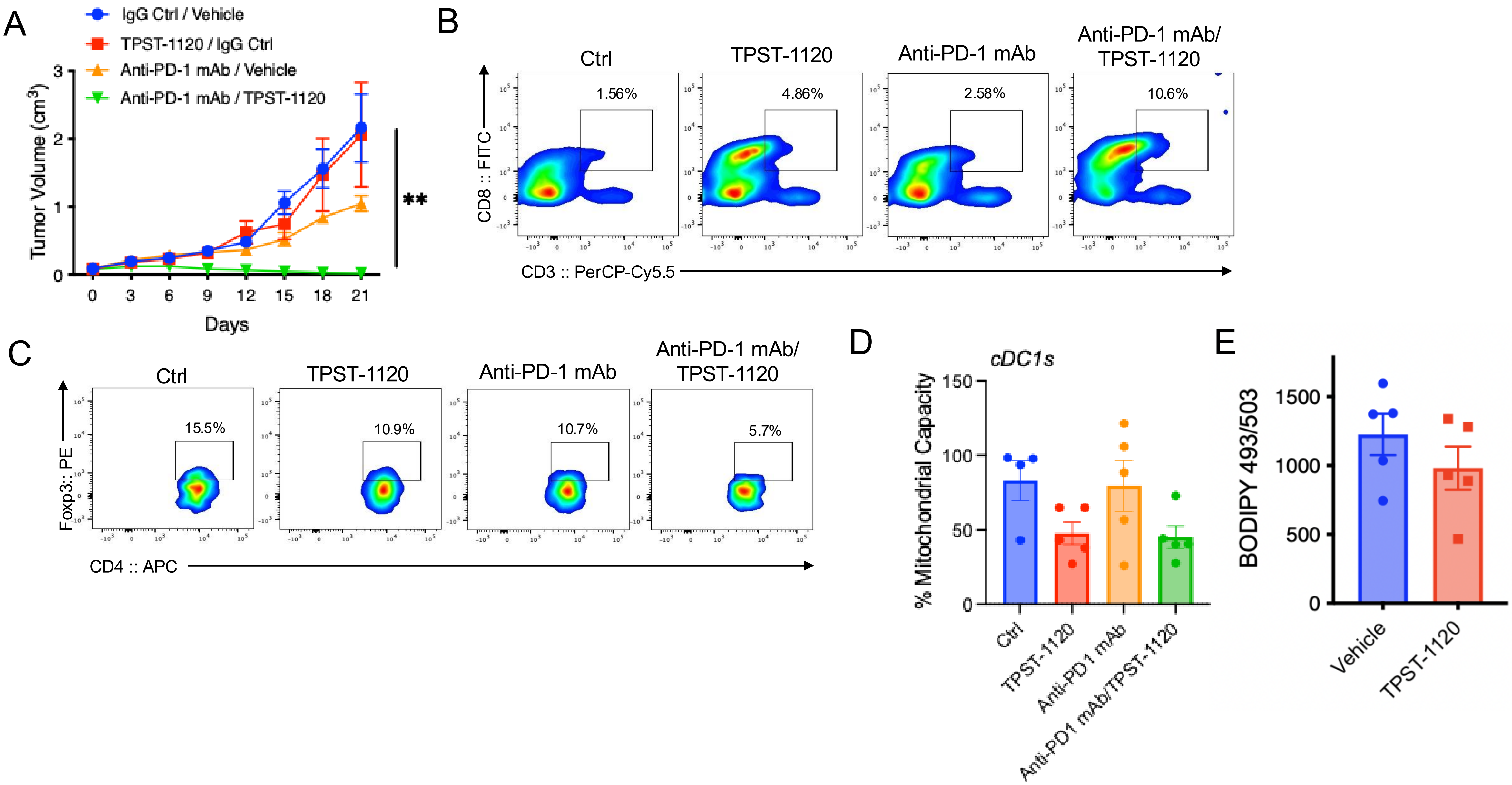
A
IgG Ctrl / Vehicle
TPST-1120 / IgG Ctrl
Anti-PD-1 mAb / Vehicle
Anti-PD-1 mAb / TPST-1120
Tumor Volume (cm³)
Days
**
B
Ctrl
TPST-1120
Anti-PD-1 mAb
Anti-PD-1 mAb/ TPST-1120
1.56%
4.86%
2.58%
10.6%
CD8 :: FITC
CD3 :: PerCP-Cy5.5
C
Ctrl
TPST-1120
Anti-PD-1 mAb
Anti-PD-1 mAb/ TPST-1120
15.5%
10.9%
10.7%
5.7%
Foxp3:: PE
CD4 :: APC
D
cDC1s
% Mitochondrial Capacity
Ctrl
TPST-1120
Anti-PD1 mAb
Anti-PD1 mAb/TPST-1120
E
BODIPY 493/503
Vehicle
TPST-1120

**Figure S7. Therapeutic Efficacy and Immune Modulation of PPARα inhibition in Combination with Anti-PD-1 Blockade. (A)** Growth curves of syngeneic $BRAF^{V600E}PTEN^{-/-}$ melanomas in mice treated with IgG isotype control/vehicle, TPST-1120 (PPAR-α antagonist), anti-PD-1 mAb/vehicle, or the combination of TPST-1120 and anti-PD-1 mAb. Representative of two independent experiments. **(B)** Representative flow cytometry plots showing the frequency of tumor-infiltrating $CD3^{+}CD8^{+}$ T cells following the indicated treatments. Representative of two independent experiments. **(C)** Representative flow cytometry plots showing the frequency of intratumoral regulatory $CD4^{+}FoxP3^{+}$ T cells following the indicated treatments. Representative of two independent experiments. **(D)** Flow cytometry assessment of mitochondrial capacity in tumor-infiltrating cDC1s across treatment groups by SCENITH assay. Data are presented as ± SEM (n=5). Representative of two independent experiments. **(E)** Quantification of neutral lipid content (BODIPY 493/503 MFI) in tumor-infiltrating cDC1s treated with vehicle or TPST-1120. Representative of two independent experiments. All data is reported as mean ± SEM (n=5). $^{**}p<0.01$ by two-way ANOVA (A). **cDC1**, type 1 conventional dendritic cell; **Ctrl**, control; **mAb**, monoclonal antibody; **MFI**, mean fluorescence intensity; **TPST**, tempest.

### Supplementary Table S1

| EV abundance | Absorbance |
|---|---|
| 3.20E+09 | 0.468199998 |
| 1.60E+09 | 0.379200011 |
| 8.00E+08 | 0.323100001 |
| 4.00E+08 | 0.295300007 |
| 2.00E+08 | 0.279500008 |
| 1.00E+08 | 0.270700008 |
| **Sample** | **Absorbance** |
| Replicate #1 | 0.322 |
| Replicate #2 | 0.292 |
| Average | 0.307 |
| EV abundance | 6.325E+08 |
| Total protein (ug) by BCA | 12.700 |
| # of exosomes per ug protein | 4.980E+07 |

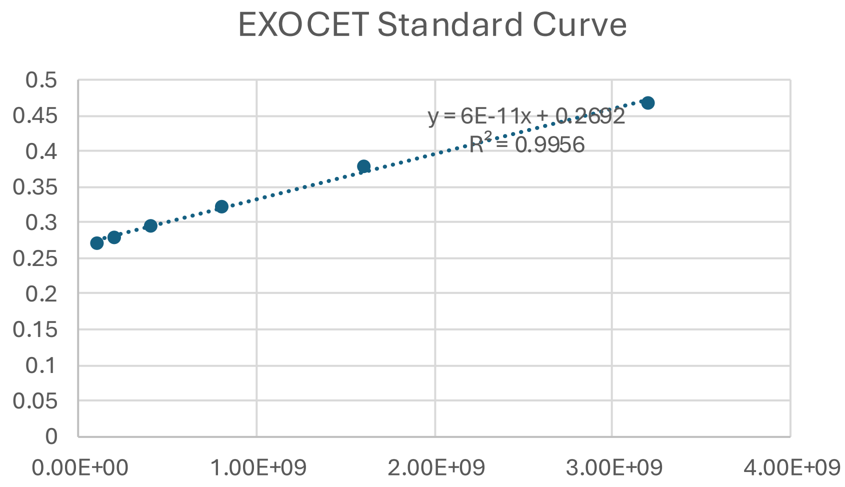


**Supplementary Table S1. EV Quantification.** Analysis performed using the EXOCET Exosome Quantification Kit. Absorbance measurements obtained using a wavelength of 405 nm (Bandwidth 9 nm, 25 flashes, 27.1C).

#### Supplementary Table S2

| Flow Cytometry Reagent | Fluorochrome | Vendor | Cat# | Clone | Dilution |
|---|---|---|---|---|---|
| LIVE/DEAD Fixable Dead Cell Stain kit (405nm) | Aqua | Thermo Fisher | L34966 | N/A | 1000 |
| LIVE/DEAD Fixable Dead Cell Stain kit (405nm) | Violet | Thermo Fisher | L34964 | N/A | 1000 |
| LIVE/DEAD Fixable Dead Cell Stain kit (488nm) | Red/PE | Thermo Fisher | 4531483 | N/A | 1000 |
| CD3 | BV421 | Biolegend | 100228 | 17A2 | 1000 |
| CD3 | PE | Biolegend | 100206 | 17A2 | 1000 |
| CD3 | PerCP/Cyanine5.5 | Biolegend | 100218 | 17A2 | 1000 |
| CD3 | FITC | Biolegend | 100204 | v | 1000 |
| CD4 | APC | Biolegend | 100412 | GK1.5 | 1000 |
| CD8 | BV421 | Biolegend | 100737 | 53-6.7 | 1000 |
| CD8 | BV510 | BD | 563068 | 53-6.7 | 1000 |
| CD8 | PE/Cyanine7 | Biolegend | 100722 | 53-6.7 | 1000 |
| CD8 | FITC | Biolegend | 100706 | 53-6.7 | 1000 |
| CD8a | PerCP/Cyanine5.5 | Biolegend | 100734 | 53-6.7 | 1000 |
| CD44 | APC | Biolegend | 103012 | IM7 | 1000 |
| CD44 | FITC | Biolegend | 156008 | NIM-R8 | 1000 |
| CD45 | APC/Cyanine7 | Biolegend | 103116 | 30-F11 | 500 |
| CD45 | PerCP/Cyanine5.5 | Biolegend | 103132 | 30-F11 | 500 |
| CD45 | Spark PLUS UV395 | LNLNDC | 103192 | 30-F11 | 500 |
| CD63 | APC | Biolegend | 143906 | NVG-2 | 1000 |
| CD63 | APC/Cyanine7 | Biolegend | 143908 | NVG-2 | 1000 |
| FOXP3 | BD Pharmingen™ PE | BD | 566881 | 3G3 | 1000 |
| F4/80 | APC | Biolegend | 123116 | NVG-2 | 1000 |
| F4/80 | FITC | Biolegend | 123108 | BM8 | 1000 |
| CD19 | APC | Biolegend | 152410 | 1D3 | 1000 |
| CD19 | FITC | BD | 553785 | 1D3 | 1000 |
| CD49b | APC | Biolegend | 103516 | HMα2 | 1000 |
| CD49b | FITC | Biolegend | 103504 | HMα2 | 1000 |
| KLRG1 | PE/Cyanine7 | Biolegend | 138416 | 2F1 | 1000 |
| CD11c | BV650 | Biolegend | 117339 | N418 | 1000 |
| CD11c | PE | Biolegend | 117308 | N418 | 500 |
| CD11c | BV510 | Biolegend | 117338 | N418 | 1000 |
| CD11c | APC | Biolegend | 117310 | N418 | 1000 |

#### Supplementary Table S2 (cont)

| Flow Cytometry Reagent | Fluorochrome | Vendor | Cat# | Clone | Dilution |
|---|---|---|---|---|---|
| I-A/I-E | APC | Biolegend | 107614 | M5/114.15.2 | 1000 |
| I-A/I-E | PE/Cyanine7 | Biolegend | 107630 | M5/114.15.2 | 1000 |
| I-A/I-E | APC/Cyanine7 | Biolegend | 107628 | M5/114.15.2 | 500 |
| XCR1 | Brilliant Violet 421 | Biolegend | 148216 | ZET | 200 |
| CD172a | PerCP/Cyanine5.5 | Biolegend | 144010 | P84 | 200 |
| CellTrace | CFSE | Thermo Fisher | C34554 | N/A | 0.74 |
| CellTrace | Violet | Thermo Fisher | C34557 | N/A | 0.74 |
| BODIPY 493/503 | N/A | Thermo Fisher | D3922 | N/A | 1.43 |
| BODIPY 581/591 C11 | N/A | Thermo Fisher | D39861 | N/A | 1.43 |
| FAO Blue | N/A | Diagnocine | FNK-FDV-0033 | N/A | 0.74 |

**Supplementary Table S2. Flow Cytometry Reagents.**

**Supplementary Table S3**

| qPCR Primers | Direction | Sequence (5' → 3') |
|---|---|---|
| *Srebf2* | Forward | GCAGCAACGGGACCATTCT |
| *Srebf2* | Reverse | CCCCATGACTAAGTCCTTCAACT |
| *Cd274* | Forward | GCTCCAAAGGACTTGTACGTG |
| *Cd274* | Reverse | TGATCTGAAGGGCAGCATTTC |
| *Pdcd1lg2* | Forward | CTGCCGATACTGAACCTGAGC |
| *Pdcd1lg2* | Reverse | GCGGTCAAAATCGCACTCC |
| *Cd200* | Forward | CTCTCCACCTACAGCCTGATT |
| *Cd200* | Reverse | AGAACATCGTAAGGATGCAGTTG |
| *Fas* | Forward | TATCAAGGAGGCCCATTTTGC |
| *Fas* | Reverse | TGTTTCCACTTCTAAACCATGCT |
| *Socs1* | Forward | CTGCGGCTTCTATTGGGGAC |
| *Socs1* | Reverse | AAAAGGCAGTCGAAGGTCTCG |
| *Xbp1s* | Forward | GCTGAGTCCGCAGCAGGT |
| *Xbp1s* | Reverse | CAGGGTCCAACTTGTCCAGAAT |
| *Sec61a* | Forward | GGAAGTCATCAAGCCATTCTGT |
| *Sec61a* | Reverse | GCATCCAGTAGAACGGGTCAG |
| *Dnajb9* | Forward | CTCCACAGTCAGTTTTCGTCTT |
| *Dnajb9* | Reverse | GGCCTTTTTGATTTGTCGCTC |

**Supplementary Table S3 (cont)**

| qPCR Primers | Direction | Sequence (5' → 3') |
|---|---|---|
| *Cpt1a* | Forward | CTCAGTGGGAGCGACTCTTCA |
| *Cpt1a* | Reverse | GGCCTCTGTGGTACACGACAA |
| *Pparα* | Forward | AGAGCCCCATCTGTCCTCTC |
| *Pparα* | Reverse | ACTGGTAGTCTGCAAAACCAAA |
| *Pparβ/δ* | Forward | TCCATCGTCAACAAAGACGGG |
| *Pparβ/δ* | Reverse | ACTTGGGCTCAATGATGTCAC |
| *Pparγ* | Forward | TCGCTGATGCACTGCCTATG |
| *Pparγ* | Reverse | GAGAGGTCCACAGAGCTGATT |
| **Genotyping Primers** | **Direction** | **Sequence (5' → 3')** |
| *Itgax-Cre* | Forward | ACTTGGCAGCTGTCTCCAAG |
| *Itgax-Cre* | Reverse | GCGAACATCTTCAGGTTCTG |
| *Srebf2* | Forward | ACCTGATGCCTTACTGTGTTACTG |
| *Srebf2* | Reverse | TCACACTTATCCCATCCGAGA |
| *Pparα Flox* | Forward | CAA GGC CAT GTC TAA TCA TCC TGG |
| *Pparα Flox* | Reverse | TCT CAT GGA TTC AAT TGA CTG ACT GG |
| *Pparα Flox* | LAR | CAA CGG GTT CTT CTG TTA GTC C |
| *Zbtb46-Cre* | WT | CTG GTT GAG GAT GAG GAT GA |
| *Zbtb46-Cre* | Mutant | GCG CTG GAG TTT CAA TAC C |

**Supplementary Table S3. Primer Sequences.**
